# Modification of xylan in secondary walls alters cell wall biosynthesis and wood formation programs

**DOI:** 10.1101/2024.05.02.592170

**Authors:** Pramod Sivan, János Urbancsok, Evgeniy N. Donev, Marta Derba-Maceluch, Félix R. Barbut, Zakiya Yassin, Madhavi L. Gandla, Madhusree Mitra, Saara E. Heinonen, Jan Šimura, Kateřina Cermanová, Michal Karady, Gerhard Scheepers, Leif J. Jönsson, Emma R. Master, Francisco Vilaplana, Ewa J. Mellerowicz

**Affiliations:** Umeå Plant Science Centre, Swedish University of Agricultural Sciences, Department of Forest Genetics and Plant Physiology, 901 83 Umeå, Sweden; RISE Research Institutes of Sweden, Drottning Kristinas väg 61, 11428 Stockholm, Sweden; Department of Chemistry, Umeå University, 901 87 Umeå, Sweden; Laboratory of Growth Regulators, Palacký University, Institute of Experimental Botany, The Czech Academy of Sciences & Faculty of Science, Olomouc, CZ-783 71 Czechia; Department of Chemical Engineering and Applied Chemistry, University of Toronto, M5S 3E5 Canada; Division of Glycoscience, Department of Chemistry, KTH Royal Institute of Technology, AlbaNova University Centre, 106 91 Stockholm, Sweden; Wallenberg Wood Science Centre (WWSC), KTH Royal Institute of Technology, 100 44 Stockholm, Sweden

**Keywords:** Glucuronoxylan, fungal xylanases, transgenic aspen, wood development, lignocellulose, secondary cell wall, hardwood genetic engineering

## Abstract

Wood of broad-leaf tree species is a valued source of renewable biomass for biorefinery and a target for genetic improvement efforts to reduce its recalcitrance. Glucuronoxylan (GX) plays a key role in recalcitrance through its interactions with cellulose and lignin. To reduce recalcitrance, we modified wood GX by expressing GH10 and GH11 endoxylanases from *Aspergillus nidulans* in hybrid aspen (*Populus tremula* L. x *tremuloides* Michx.) and targeting the enzymes to cell wall. The xylanases reduced tree height, modified cambial activity by increasing phloem and reducing xylem production, and reduced secondary wall deposition. Xylan molecular weight was decreased, and the spacing between acetyl and MeGlcA side chains was reduced in transgenic lines. The transgenic trees produced hypolignified xylem having thin secondary walls and deformed vessels. Glucose yields of enzymatic saccharification without pretreatment almost doubled indicating decreased recalcitrance. The transcriptomics, hormonomics and metabolomics data provided evidence for activation of cytokinin and ethylene signaling pathways, decrease in ABA levels, transcriptional suppression of lignification and a subset of secondary wall biosynthetic program, including xylan glucuronidation and acetylation machinery. Several candidate genes for perception of impairment in xylan integrity were detected. These candidates could provide a new target for uncoupling negative growth effects from reduced recalcitrance. In conclusion, our study supports the hypothesis that xylan modification generates intrinsic signals and evokes novel pathways regulating tree growth and secondary wall biosynthesis.

## Introduction

Plant cell wall is a highly dynamic and heterogenous structure made by complex chemical organization of cellulose, diverse matrix polysaccharides, structural proteins and polyphenols (Albersheim et al., 2010). The structure, composition and molecular interaction of matrix polysaccharides determine cell shape and tensile properties necessary for the mechanical strength of cell wall. Xyloglucans and pectins form the matrix of primary cell wall and their interactions with cellulose microfibrils within highly hydrated architecture facilitate cell expansion. The secondary walls (SWs) are deposited in xylem cells after cell expansion and have a denser and thicker network of cellulose microfibrils with a SW-specific combination of matrix polysaccharides including glucuronoxylan (GX) and glucomannan. Deposition of this SW polysaccharide network and its subsequent lignification starting from the primary wall makes further cell expansion impossible but provides xylem cells with mechanical strength and rigidity. The primary and secondary walls constitute wood biomass which is the most abundant renewable resource on Earth for sustainable production of eco-friendly materials, chemicals and energy carriers (Keegan et al. 2013; Bar-On et al. 2018; Martínez-Abad et al. 2018).

Biosynthesis of cell wall components has been largely investigated by studying cell wall mutants like *murus* (*mur)* (Mertz et al., 2012), *irregular xylem* (*irx*) (Turner and Somerville, 1997), *fragile stem* (*fra*) (Zhong et al., 2005), *trichome birefringence-like* (*tbl*) (Potikha and Delmer, 1995) and by systematic gene sequence analyses (Cantarel et al., 2009; Kumar et al., 2019). The enzymatic activities of several proteins have been characterized (e.g. Cavalier and Keegstra 2006; Maris et al. 2011). However, it is still not well understood how the different activities are coordinated during cell wall biosynthesis to produce cell walls of required properties. Knowledge on the dynamic macromolecular changes during cell wall formation and modification in response to developmental and environmental changes is fundamental for our efforts to create plants with desired cell wall chemical composition suitable for industrial applications such as biorefinery and production of biomaterials (Somerville and Bonetta 2001; Pauly and Keegstra, 2008, 2010).

The secondary cell wall of hardwood xylem contains approx. 25% (dry weight) of GX. GX has been reported to have distinct structural variants in terms of relative abundance of acetylation and glucuronidation and the pattern of these decorations resulting in the major xylan domain that forms a two-fold helical screw conformation, forming compatible region for hydrogen bonding to the hydrophilic surface of cellulose, and a minor domain forming a three-fold screw making regions interacting with lignin (Bromley et al., 2013; Busse-Wicher et al., 2014; Simmons et al., 2016; Yuan et al., 2016a; 2016b; 2016c; Grantham et al., 2017). The unsubstituted xylan surface can form stacking interactions on the hydrophobic surface of cellulose (Gupta et al., 2021). Thus, xylan molecules can differently affect cell wall architecture, and how their biosynthetic process is controlled to ensure formation of functional cell wall is not understood. This is complicated by the fact that in addition to enzymatically driven xylan biosynthesis and modification there are spontaneous processes that may occur in the cell wall changing its properties. For example, there is a long-standing hypothesis that xylan provides nucleation sites for lignin polymerization. In grasses, the ferulic acid linked to arabinose side chains of xylan is believed to initiate lignification (Markwalder & Neukom, 1976; Hartley et al., 1990; Ralph et al., 1995). In poplar, Ruel et al (2006) proposed that hemicellulose-lignin covalent linkage serves as an anchor for lignin polymerization. The extracellular lignin analysis in spruce cell cultures suggests that exogenously supplied xylan can act as a nucleation center for lignin polymerization, depending on the amount and solubility of the xylan provided (Sapouna et al., 2023).

Postsynthetic modification of the cell wall by overexpression of xylan-modifying microbial enzymes represents a promising strategy to examine the contributions of different xylan structures to cell wall functions and to investigate mechanisms regulating cell wall biosynthesis in response to xylan integrity defects (Pogorelko et al., 2011). The cell wall modification by overexpression of xylan-acting microbial enzymes has also been demonstrated to decrease biomass recalcitrance (Pogorelko et al., 2011; Gandla et al., 2015; Pawar et al., 2016; 2017; Pramod et al., 2021). Similarly, xylan structure defects caused by suppression of native xylan biosynthetic genes altered plant cell wall architecture and improved saccharification (Donev et al., 2018). These experiments demonstrated the potential use of plants compromised in xylan integrity for biorefinery applications, and in some cases revealed activation of biotic stress and growth responses triggered by xylan modification. However, their effects on cell wall developmental pathway received little or no attention.

In the present study, we report changes in xylem cell wall chemistry and resulting modifications in cell wall biosynthesis and xylem cell developmental programs in transgenic aspen overexpressing endo-1,4-β-D-xylanases of GH10 and GH11 families from *Aspergillus nidulans* in apoplast of developing xylem cells. Endo-1,4-β-D-xylanases (EC 3.2.1.8) cleave internal 1,4-β-xylosidic bonds in xylan backbones, producing low molecular weight (MW) heteroxylans and unsubstituted or branched xylo-oligosaccharides (XOS) (Reilly, 1981; Pollet et al., 2010). The products of GH10 and GH11 xylanases sightly differ because only GH10 can accommodate a substituted xylosyl residue at the −1 subsite of the active site whereas both families require unsubstituted xylosyl residue at the +1 subsite (Biely et al., 1997; Pell et al., 2004; Kolenová et al., 2006; Vardakou et al., 2008; Kojima et al., 2022). The expression of xylanases altered xylem cell wall biosynthetic program and modified cambial activity suggesting the loss of xylan integrity in SWs is sensed by differentiating xylem cells. The resulting lignocellulosic biomass had substantially increased saccharification potential. However the plants’ growth was affected and uncoupling of the two effects is needed before such a strategy could be used for practical deployment.

## Results

### Microbial xylanases affected growth and vascular tissue differentiation pattern in aspen

Transgenic aspen expressing fungal xylanases showed clear morphological changes (**Fig. 1A**). All growth parameters (stem height and diameter, aboveground and root biomass) were significantly affected compared to the wild-type (WT) (**Fig. 1B**). Transgene transcript levels were higher in 35S promoter lines than in WP lines (**Fig. S1**), which did not correlate with growth penalty. However, in three WP:GH10 lines with different transgene levels, there was a clear negative impact of transgene transcript level on height and biomass production. Although radial growth was not affected in the majority of transgenic lines, the measurement of secondary vascular tissues from transverse sections revealed increased secondary phloem and decreased secondary xylem production (**Fig. 1C**). The pith area of transgenic lines was also significantly increased. These observations indicate that xylanases stimulated growth of pith and had a major impact on cambial activity shifting it from xylem to phloem production.

**Figure 1.**
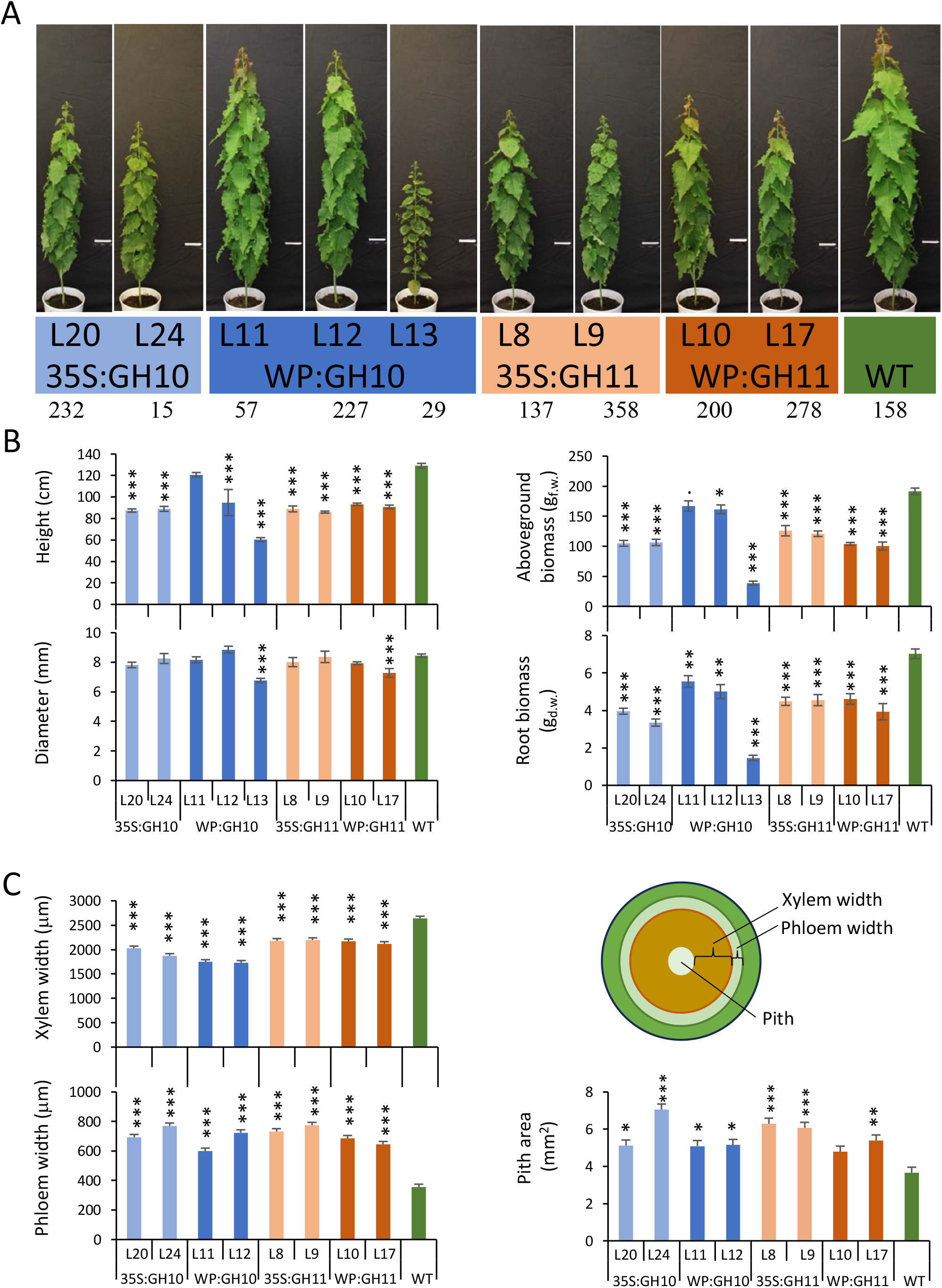
Growth of transgenic lines expressing GH10 and GH11 xylanases. **(A)** Morphology of 9-week-old plants. Size bar =10 cm. **(B)** Plant size. **(C)** The width of secondary xylem and secondary phloem and the area of pith measured in stem cross sections of internode 40, as shown on the diagram. Data are means ± SE; N=6 trees for transgenic lines and 14 for WT for (B), 2 trees x 10 radii or 2 trees x 2 sections in (C). * - P≤0.05; ** - P≤0.01; *** - P≤0.001 for comparisons with WT by Dunnett’s test.

Intriguingly, the appearance of freshly cut stems of transgenic lines was altered. All lines, but the low-expressing line WP:GH10_11, showed a markedly increased zone of wet xylem, which normally indicates developing not fully lignified xylem (**Fig. 2A**). The stems were also much easier to cut, suggesting changes in cell wall properties. Indeed, SilviScan analysis (**Fig. 2B**) showed that several transgenic lines had higher wood density or increased cellulose microfibril angle (MFA). The number of xylem cells per radial file was reduced in transgenic lines confirming microscopy analyses. Furthermore, an increase in vessel fraction with concomitant decrease in vessel perimeter and an increase in fiber diameter were observed in several transgenic lines, suggesting that xylem cell fate and xylem cell expansion were also affected by the xylanases.

**Figure 2.**
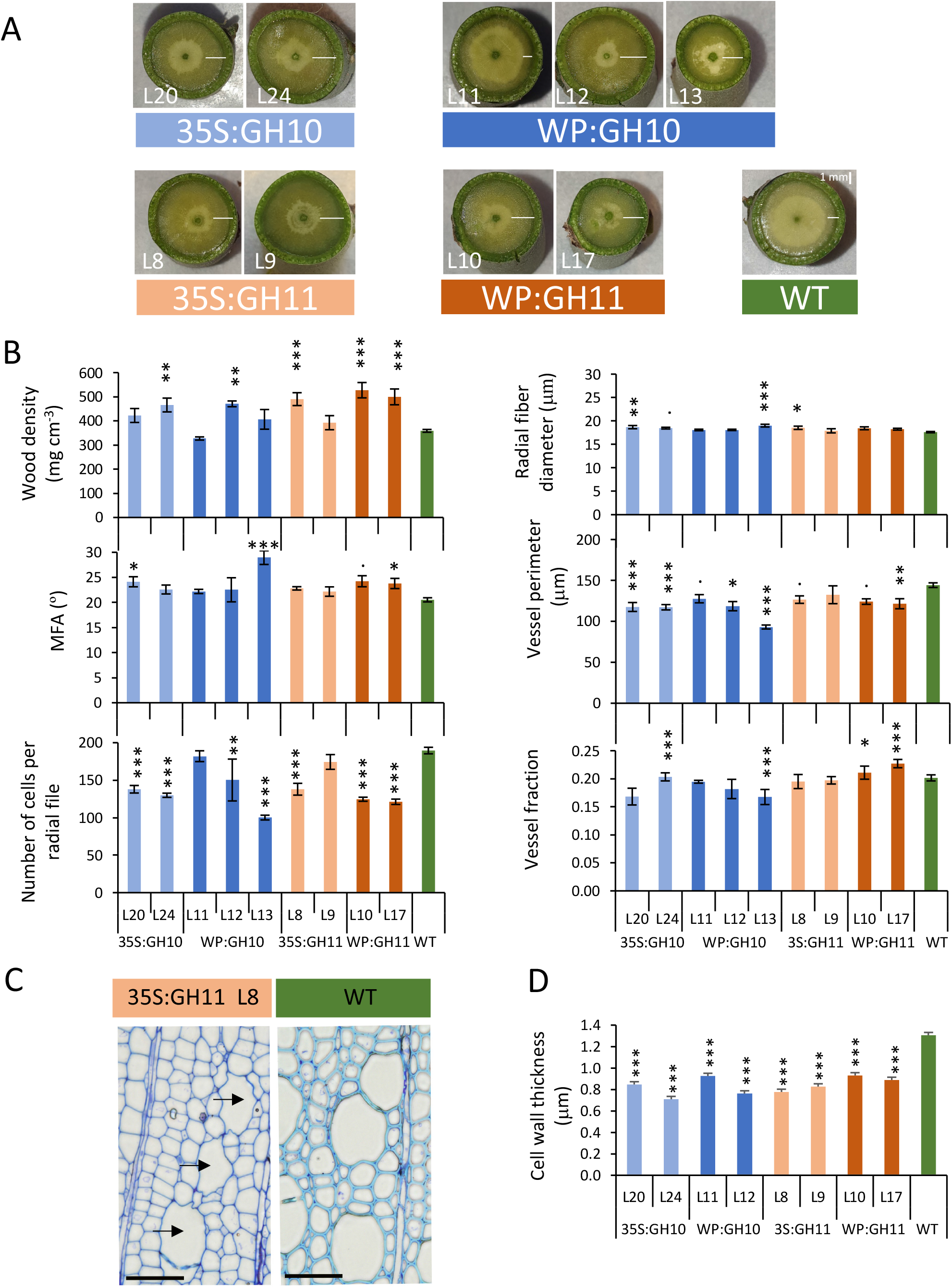
Wood quality traits of transgenic lines expressing GH10 and GH11 xylanases determined by SilviScan and anatomical analyses. **(A)** Appearance of SilviScan wood samples freshly dissected from the stems. Note the wider wet-looking zone (white bars) in transgenic lines. **(B)** Different wood quality traits measured by SilviScan. **(C)** Typical appearance of wood in xylanase-expressing plants. Note a reduction in cell wall thickness*, irregular xylem* phenotype (collapsed vessels, black arrows) and altered cell wall staining properties. Toluidine blue stained wood cross sections. Sections of other lines are shown in Supplementary Figure S2. **(D)** Secondary wall thickness measured by transmission electron microscopy analysis. WT-wild type, MFA – cellulose microfibril angle. Data in B and D are means ± SE, N = 6 for transgenic lines and 24 for WT in B, or 2 trees x 3 images x 4 measurement in D. * - P≤0.05; ** - P≤0.01; *** - P≤0.001 for comparisons with WT by Dunnett’s test.

Analysis of semi-thin transverse sections stained with toluidine blue O (TBO) revealed a substantial decrease in cell wall thickness and frequent occurrence of *irregular xylem* (*irx*) phenotype (**Fig. 2CD, Fig. S2**). Moreover, a shift in TBO color from cyan in WT to violet-blue in transgenic lines suggested decrease in lignification. This was confirmed by analysis of lignin autofluorescence in the wood sections of transgenic and WT plants (**Fig. S3**). Lignin autofluorescence images also revealed large wood areas in transgenic plants with very low signal, which possibly represent patches of tension wood (TW). All these changes were attenuated in the line WP:GH10_11 that had lower transgene expression compared to other lines.

### Xylanase expression had a major impact on the content and composition of wood matrix sugars and lignin

Sulfuric acid hydrolysis showed no consistent changes in cellulose (glucan) content (**Fig. 3A**). On the other hand, acid methanolysis-TMS analysis showed significant changes in matrix sugars (**Fig. 3B**): xylose, MeGlcA and GlcA contents decreased in most or all transgenic lines, mannose contents decreased in most lines but WP:GH10 and most lines had lower glucose unit content than WT. WP:GH10 lines showed increase in pectin-related sugars including rhamnose, galacturonic acid, galactose and arabinose, whereas the opposite trend or no change was observed for other lines. Wood analysis by Py-GC/MS revealed a significant decrease in total lignin and guaiacyl (G) unit contents in transgenic lines (**Fig. 3C**). Whereas the G-lignin units were substantially reduced in all transgenic lines, the syringil (S) lignin units were decreased only in GH11-expressing lines. The content of other phenolics was on the other hand increased in the majority of the transgenic lines. Spatial distribution pattern of lignin and xylan in cell walls analyzed by transmission electron microscopy revealed a severe depletion of lignin in the compound middle lamellae and SW layers of xylem fibers in transgenic trees (**Fig. 4A**) and a significant decrease in gold particle density labeling LM10 xylan epitopes in transgenic lines (**Fig. 4BC**). Thus, cell wall analyses revealed major impact of xylanases on the lignin and xylan in wood cell walls.

**Figure 3.**
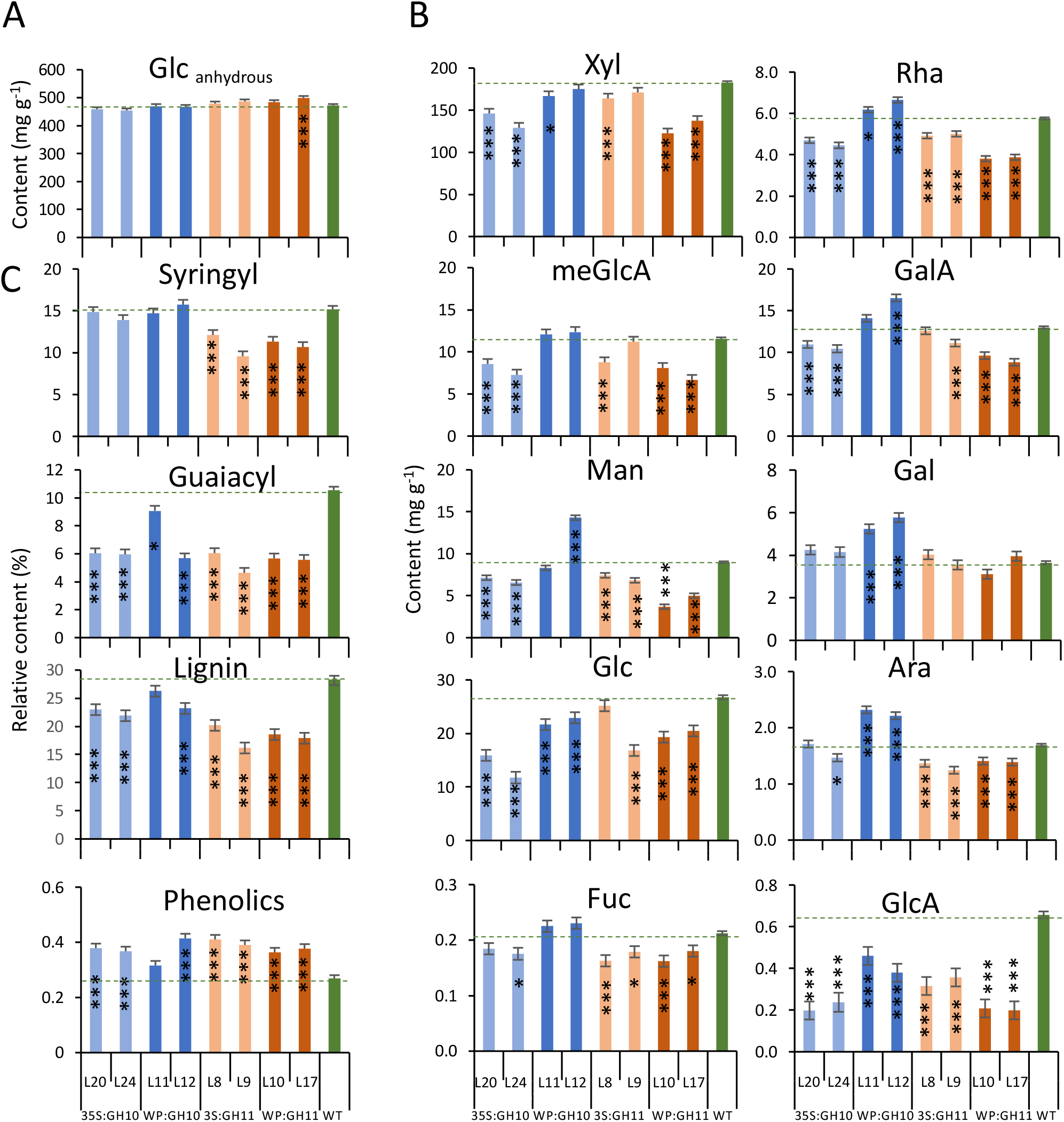
Chemical composition of wood in transgenic lines expressing GH10 and GH11 xylanases. Glucan (anhydrous glucose) content in dry wood determined by sulfuric acid hydrolysis **(A)**. Matrix sugar content (hydrous) determined by methanolysis-TMS per dry weight of dry destarched alcohol-insoluble wood **(B).** Relative content of syringyl (S) and guaiacyl (G) monolignols, total lignin (S+G+H) and phenolics in wood powder determined by the pyrolysis GC-MS **(C).** H - *p*-hydroxyphenyl units. Data are means ± SE, N = 3 technical replicates of pooled material from 6 trees in A; N = 9 (3 technical and 3 biological replicates in B: N = 3 biological replicates for C., * - P≤0.05; ** - P≤0.01; *** - P≤0.001 for comparisons with WT by Dunnett’s test.

**Figure 4.**
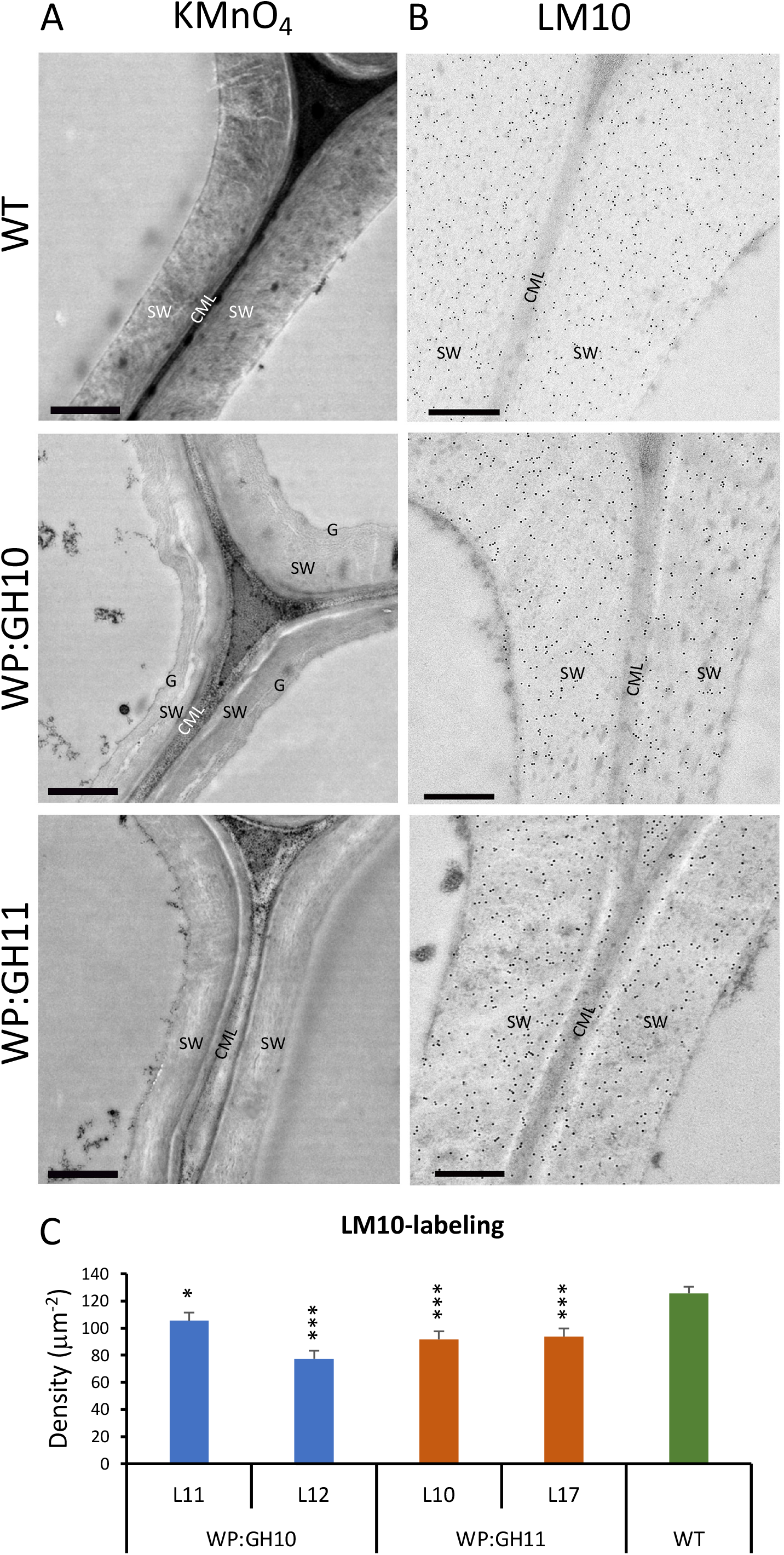
Transmission electron microscopy of cell walls in the xylem of transgenic lines expressing GH10 and GH11 xylanases showing differences in lignin and xylan content compared to wild type (WT). **(A)** Lignin in the fiber walls detected with KMnO_4_ which is seen as a dark deposit in the compound middle lamella (CML) and secondary walls (SW) is highly reduced in transgenic lines. Note also the presence of G-layer (G) in one of the transgenic samples. **(B, C)** Immunogold localization of xylan in fiber cell walls using LM10 antibody **(B)** and quantification of gold particle density over secondary walls **(C)**. Scale bar =1 µm in A and 500 nm in B; data in C are means ± SE, N = 2 trees x 3 images x 4 measurements. * - P≤0.05; ** - P≤0.01; *** - P≤0.001 for comparisons with WT by Dunnett’s test.

### Detailed xylan analysis revealed that xylanases affected its molecular structure

The yield of xylan in 20 and 30 min subcritical water extracts (SWE) determined as sum of xylose and MeGlcA contents was increased in WP:GH11 lines indicating increased xylan solubility compared to WT (**Fig. 5A**). The xylan from 30 min extracts of transgenic lines was characterized by a higher degree of acetylation (**Fig. 5B**). The molar mass distribution of 30 min SWE determined by size exclusion chromatography revealed a decrease in molecular weight in transgenic trees indicating the reduction in the degree of polymerization of xylan according to the expected cleavage activity of expressed xylanases (**Fig. 5C**).

**Figure 5.**
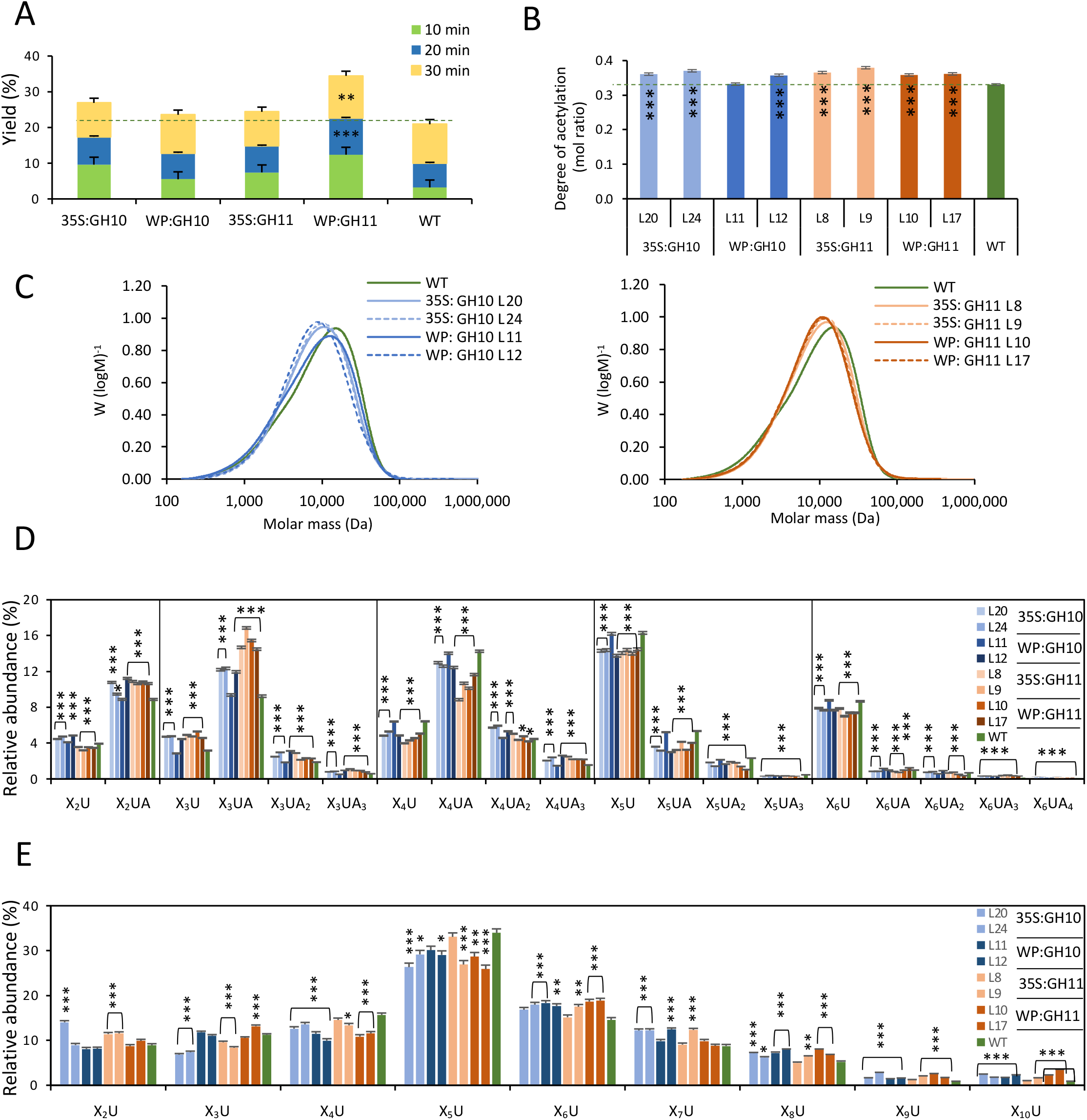
Characterization of xylan structure in transgenic lines expressing GH10 and GH11 xylanases. **(A)** Xylan yields during successive steps of subcritical water extraction relative to starting xylan weight. **(B)** The degree of acetylation of xylan extracted by SWE for 30 min. **(C)** Size exclusion chromatography of 30 min SWE extract. **(D, E)** Oligomeric mass profiling (OLIMP) by ESI-MS of SWE-extracted glucuronoxylan **(D)** or alkali-extracted glucuronoxylan **(E)** hydrolyzed with GH30 glucuronoxylanase. Relative abundance of oligosaccharides in D and E are calculated from the total ESI-MS intensities. Data in A, B, D, E are means ± SE, N= 2 lines in A, 2 technical replicates in B, and 3 technical replicates in D and E, * - P≤0.05; ** - P≤0.01; *** - P≤0.001 for comparisons with WT by Dunnett’s test.

The oligomeric mass profiling (OLIMP) of acetylated xylan from SWE and digested with GH30 glucuronoxylanase showed an increased population of oligomers representing closer glucuronidation spacing and higher acetyl substitution (X_2_UA, X_3_UA) and a decreased abundance of oligomers representing more spaced substitutions (X_4_UA, X_5_U, X_6_U) in transgenic lines (**Fig. 5D, S4**). OLIMP of alkali-extracted xylan showed in opposite a significant decrease in oligomers representing closer glucuronidation (X_3_U to X_5_U) and increase in those representing more spaced glucuronidation (X_6_U to X_10_U) (**Fig. 5E, S5**). Altogether these data indicate that the xylan domains with close MeGlcA and acetyl substitution are protected from GH10 and GH11 xylanases expressed in transgenic lines, and that regions with highly spaced glucuronidation are most likely hindered from GH10 and GH11 xylanases by high acetyl substitution.

### Xylanases-induced cell wall chemical changes improved saccharification potential of wood

Wood of xylanase-expressing lines showed substantial reduction of recalcitrance which was particularly evident in saccharification without pretreatment. Glucose production rate, and glucose and xylose yields increased up to 210%, 190% and 300% of WT levels, respectively (**Fig. 6A**). The improvements for GH11 were more substantial compared to GH10 (even when disregarding line WP:GH10_11), as supported by P_contrast GH10_ *_vs_* _GH11_ ≤0.0001 for all three parameters. After pretreatment, glucose production rates were also increased in transgenic lines, but to a lesser extent (up to 130%), whereas only GH11 lines showed higher glucose yields (up to 130%) than WT (**Fig. 6B**). Total xylose yields were increased for some lines (35s:GH11) but reduced for others (35s:GH10 and WP:GH11) reflecting the net decrease in xylose unit content of these lines. Furthermore, the yields of mannan and galactan in pretreatment liquid were altered in many transgenic samples reflecting changes in their content and solubility (**Fig. S6**).

**Figure 6.**
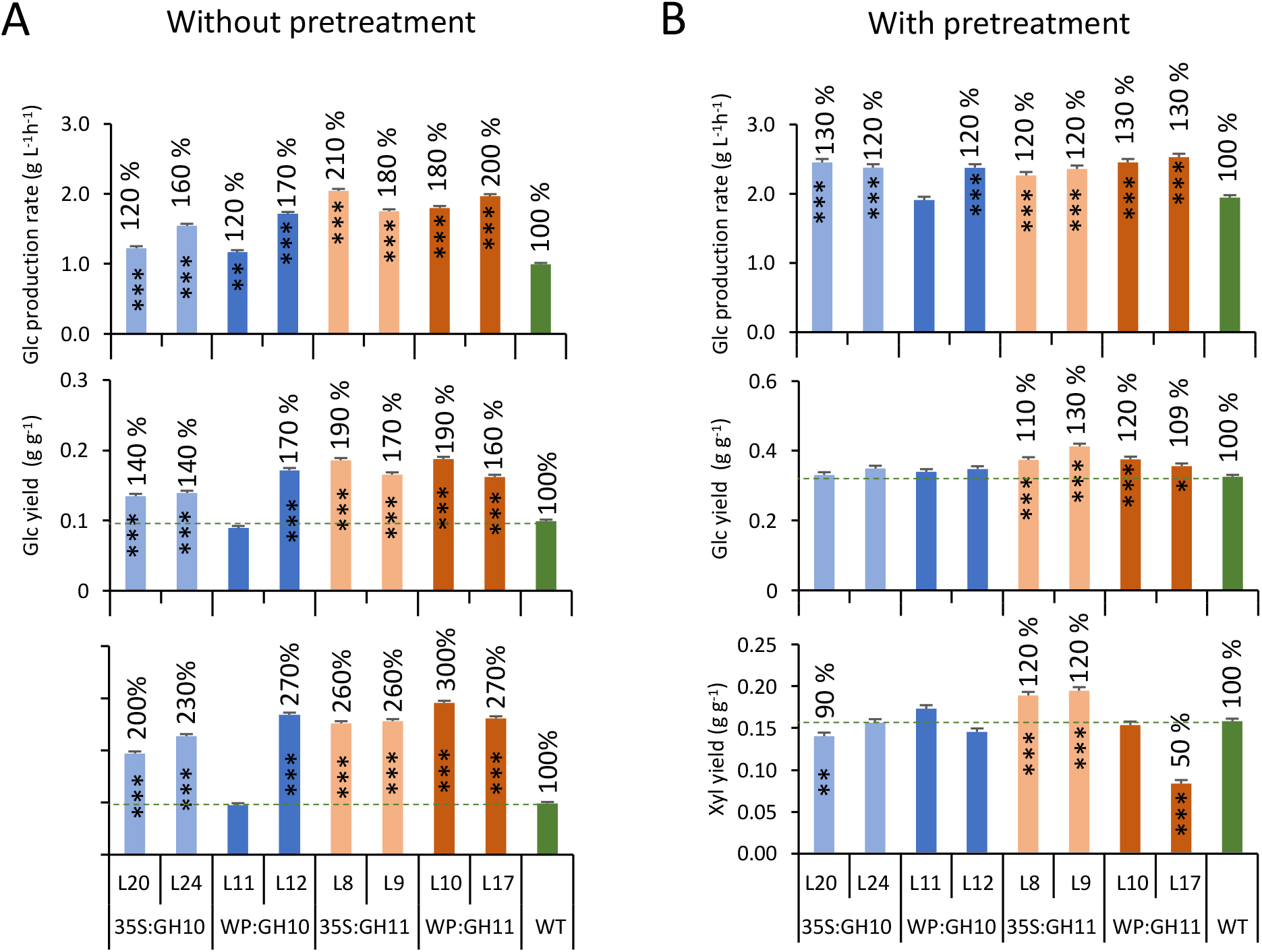
Effects of *in planta* expression of GH10 and GH11 xylanases on saccharification of wood. Glucose production rates, and glucose and xylose yields in saccharification without **(A)** and with **(B)** acid pretreatment. Data are means ± SE, N = 3 or 6 technical replicates from the pooled material of 6 trees for transgenic lines and WT, respectively. * - P≤0.05; ** - P≤0.01; *** - P≤0.001 for comparisons with WT by Dunnett’s test.

### Changes in hormonomics and metabolomics provide evidence for intrinsic regulation of vascular differentiation and cell wall lignification in transgenic trees

To understand mechanism of developmental changes triggered by xylan integrity impairments in SWs, we analyzed hormones in developing wood of WP:GH10 and WP:GH11 lines and WT. There was an overall similarity in hormonal changes induced by GH10 and GH11, with many cytokinin forms, some auxin forms, abscisic acid (ABA) and the ethylene precursor 1-aminocyclopropane-1-carboxylic acid (ACC) being significantly affected, whereas no changes were seen in jasmonates (JA) or salicylic acid (**Fig. 7A**). Significant increases in active forms of cytokinins, trans-zeatin and N^6^-isopentenyladenine, and their riboside precursors were evident indicating elevated cytokinin signaling (**Fig. 7B**). In contrast, the levels of indole-3-acetic acid (IAA) were decreased with concomitant increases in inactivated IAA forms. The ABA showed a two-fold decrease while ACC concentration increased almost four times in transgenic trees, indicating altered stress signaling *via* ABA and ethylene. This provides evidence that hormones regulating the cambial activity, xylem differentiation and stress responses were affected by expression of xylanases in xylem cells forming SWs.

**Figure 7.**
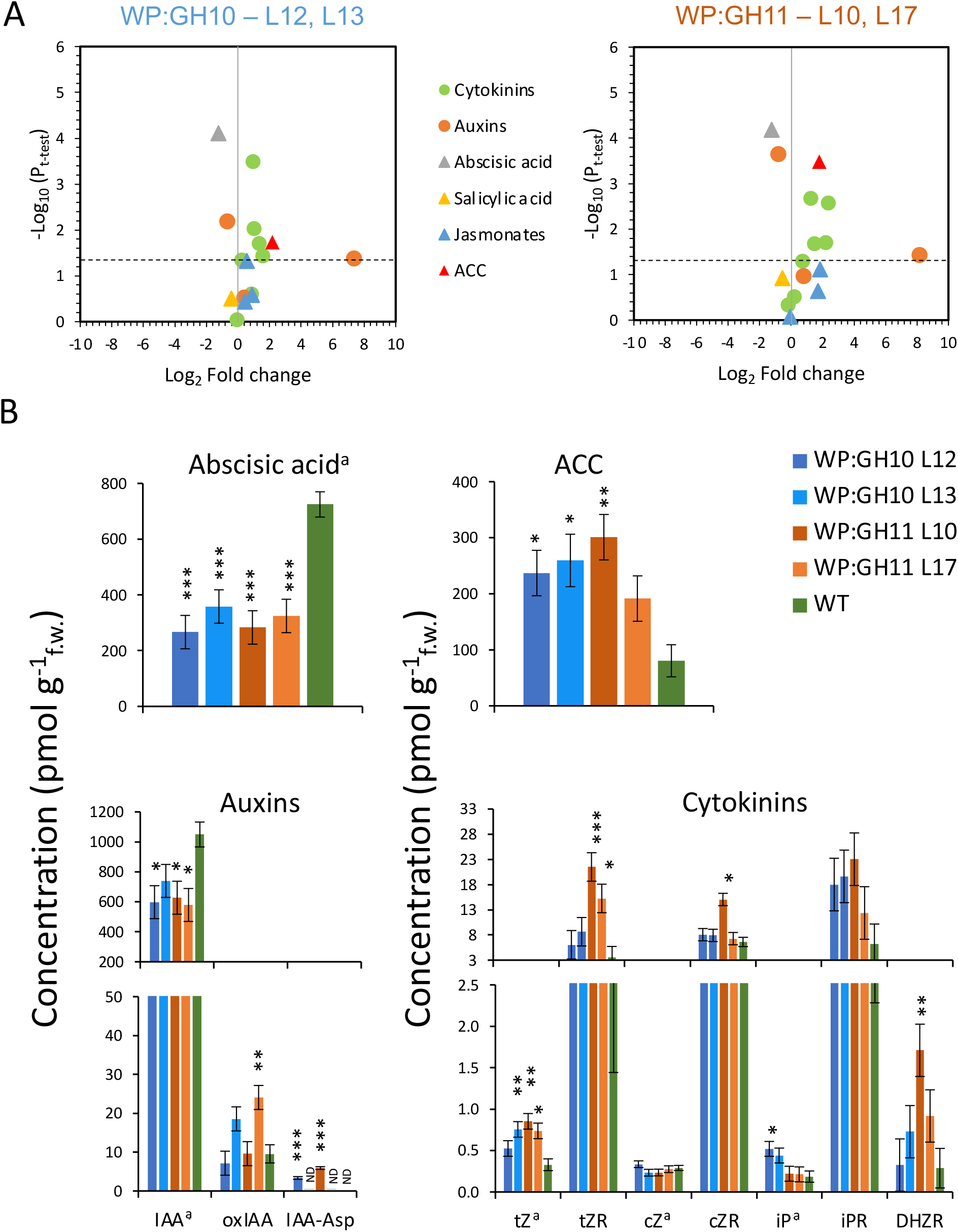
Changes in hormonal status in wood forming tissues of transgenic lines expressing GH10 or GH11 xylanases. **(A)** Volcano plots showing all detected hormones. Colored signs above the dashed lines show the hormones that were either significantly increased or reduced in the transgenic lines at P≤0.05 compared with wild type (WT). **(B)** Bar plots showing mean contents of abscisic acid, ACC, auxins and cytokinins in transgenic lines as compared to WT. Data are means ± SE, N = 4 trees for transgenic lines and 7 for WT; * - P≤0.05; ** - P≤0.01; *** - P≤0.001 for comparisons with WT by Dunnett’s test. ND – not detected; active hormones are marked with “a”. ACC - 1-aminocyclopropane-1-carboxylic acid; IAA – indole-3-acetic acid; oxIAA - 2-oxoindole-3-acetic acid; IAA-Asp – IAA-aspartate; *t*Z – *trans*-zeatin; *t*ZR – *trans*-zeatin riboside; *c*Z – *cis*-zeatin; *c*ZR – *cis*-zeatin riboside; iP - N^6^-isopentenyladenine; iPR - N^6^-isopentenyladenosine; DHZR – dihydrozeatin riboside.

Xylanases induced striking changes in the metabolomes of transgenic lines, which were highly similar between WP:GH10 and WP:GH11 lines as shown by the volcano plots and Venn diagrams (**Fig. 8AB**). Significantly affected compounds detected by GC-MS analysis were mostly upregulated. They comprised amino acids and sugars, including xylose (Xyl) and xylobiose (Xyl-B) (**Fig. 8C**). LC-MS analysis revealed metabolites mostly reduced in transgenic lines of which the most affected were lignols (some with over 30-fold decrease), phenolic glycosides and phenyl propanoid-related metabolites (**Fig. 8A-C**), demonstrating the specific impact on lignin biosynthetic pathway.

**Figure 8.**
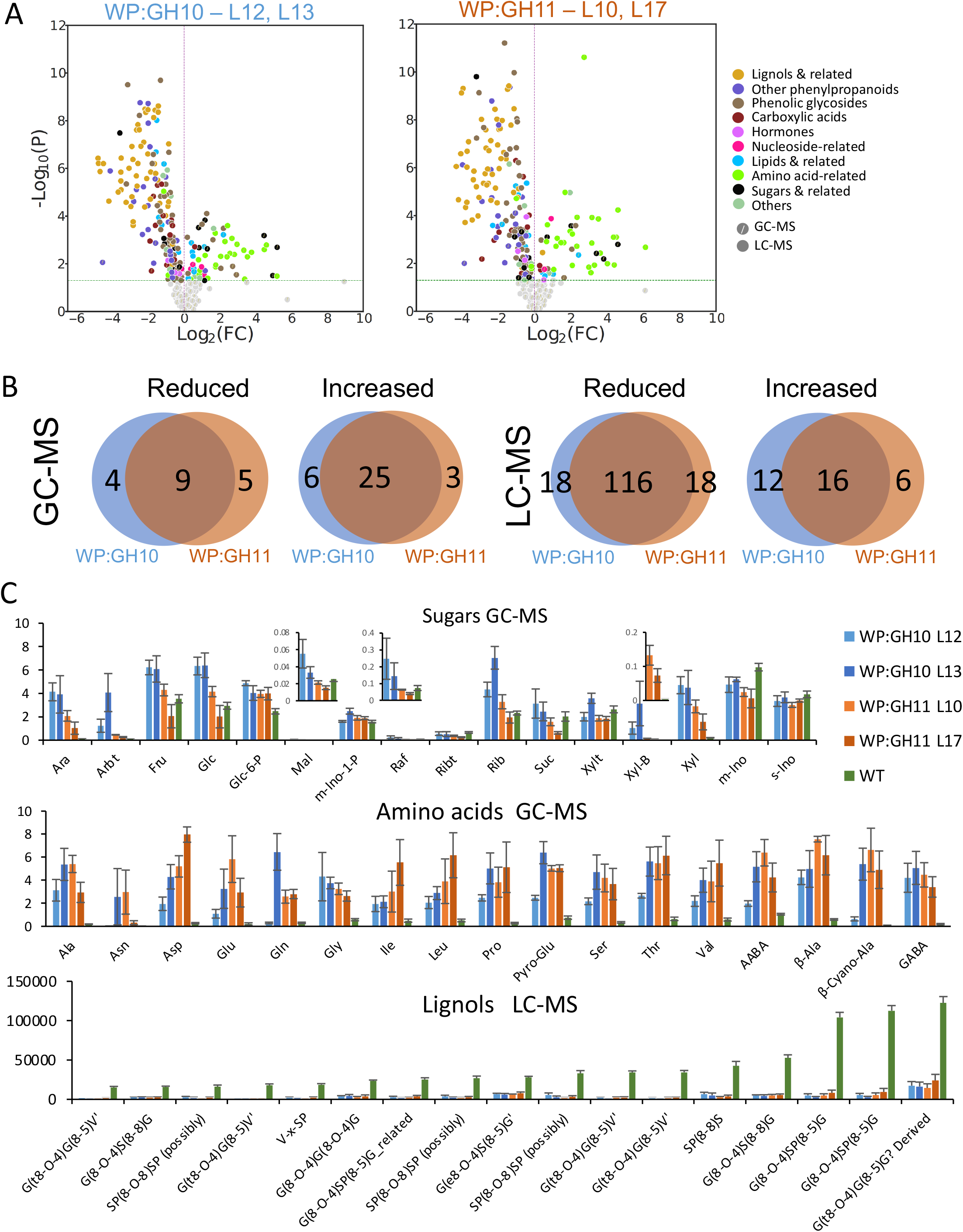
Metabolomes of developing wood in transgenic lines expressing xylanases show massive changes in several groups of compounds. **(A)** Volcano plots of metabolites analyzed by LC-MS and GC-MS showing groups of compounds significantly affected (P≤0.05, t-test) in transgenic lines compared to wild type (WT). **(B)** Venn diagrams showing number of metabolites significantly affected transgenic lines compared to WT. **(C)** Quantitative variation in integrated peaks (in relative units) corresponding to the most affected groups of compounds (amino acids, sugars and most abundant lignols). Data are means ± SE, N= 8 trees for WT and 4 for transgenic lines. Complete lists of metabolites are shown in Tables S1 – S3.

### Transcriptomic changes

#### Overview of transcriptomic changes in WP:GH10 and WP:GH11 transgenic lines

RNA-seq analysis of developing xylem identified 1600 - 2700 differentially expressed genes (DEG) in WP:GH10 and WP:GH11 lines, with an overrepresentation of upregulated genes (**Fig. 9A, Supplementary Table S4**). The core genes affected in common for all xylanase-expressing lines included 391 upregulated and 239 downregulated genes (**Supplementary Table S5**). Gene ontology (GO) enrichment analysis of these genes revealed upregulation in GO terms related to the photosynthesis, chlorophyll binding, generation of energy and stress and downregulation in categories related to lipid, protein and amino acid metabolism, oxidoreductase activities, and cell wall biosynthesis (**Fig. 9B, Supplementary Table S6**).

**Figure 9.**
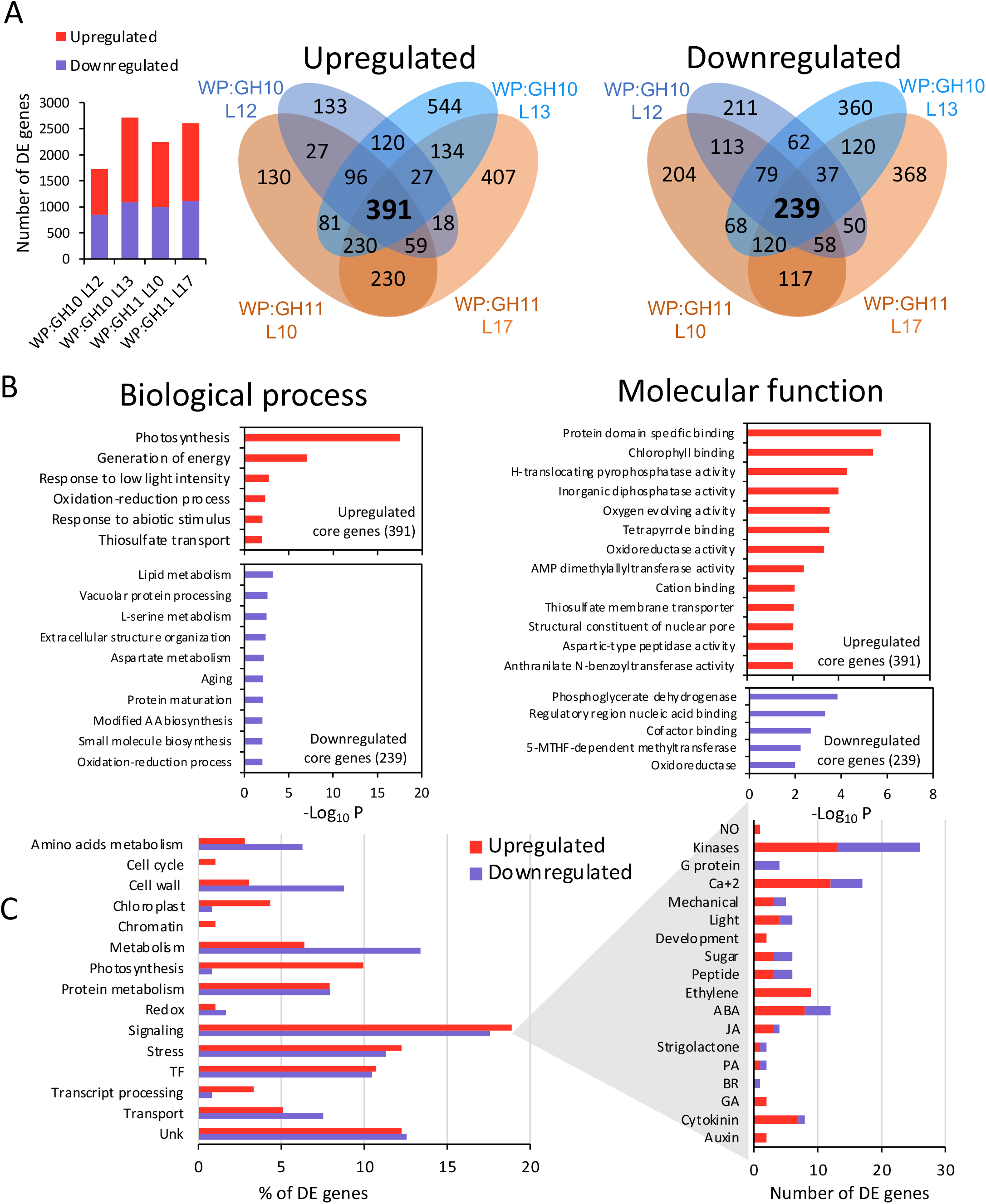
Expression of GH10 and GH11 xylanases alters transcriptomes of wood forming tissues in transgenic lines. **(A)** Numbers of differentially expressed (P_adj_≤0.01, |Log_2_(fold change)|≤0.58) genes and associated Venn diagrams. Number of core genes differentially expressed in both transgenic lines of each construct are shown in bold. **(B)** Gene ontology (GO) enrichment analysis for the core DE genes. **(C)** DE genes by different functions to which the genes were exclusively assigned. The functional classification is listed in Table S7. 5-MTHF – 5-methyltetrahydrofolate; AA-amino acid; ABA – abscisic acid; BR – brassinosteroids; GA – gibberellins; JA – jasmonate; NO – nitric oxide; PA – polyamines; TF – transcription factor.

The exclusive functional classification of the core genes (**Fig. 9C, Supplementary Table S7**) showed that signaling, stress and transcription factors functions were most highly represented among both up-and downregulated genes. Interestingly, many of the stress-related genes were annotated as responsive to anoxia. “Amino acid metabolism” and “cell wall” categories were highly represented among the downregulated genes whereas a. This indicates that changes seen in about 10% of upregulated genes were associated with photosynthesis.

#### Transcriptomic changes in signaling and stress response genes

Since the signaling was the most represented function among DEGs, we analyzed these genes in more detail. Transcripts for different kinases and calcium signaling genes were the two most highly represented groups in the signaling category (**Fig. 9C**, **Table 1**). Of the hormone-related genes, those related to ABA, ethylene, and cytokinins were most highly represented, which is in good agreement with the hormone analyses.

**Table 1.**
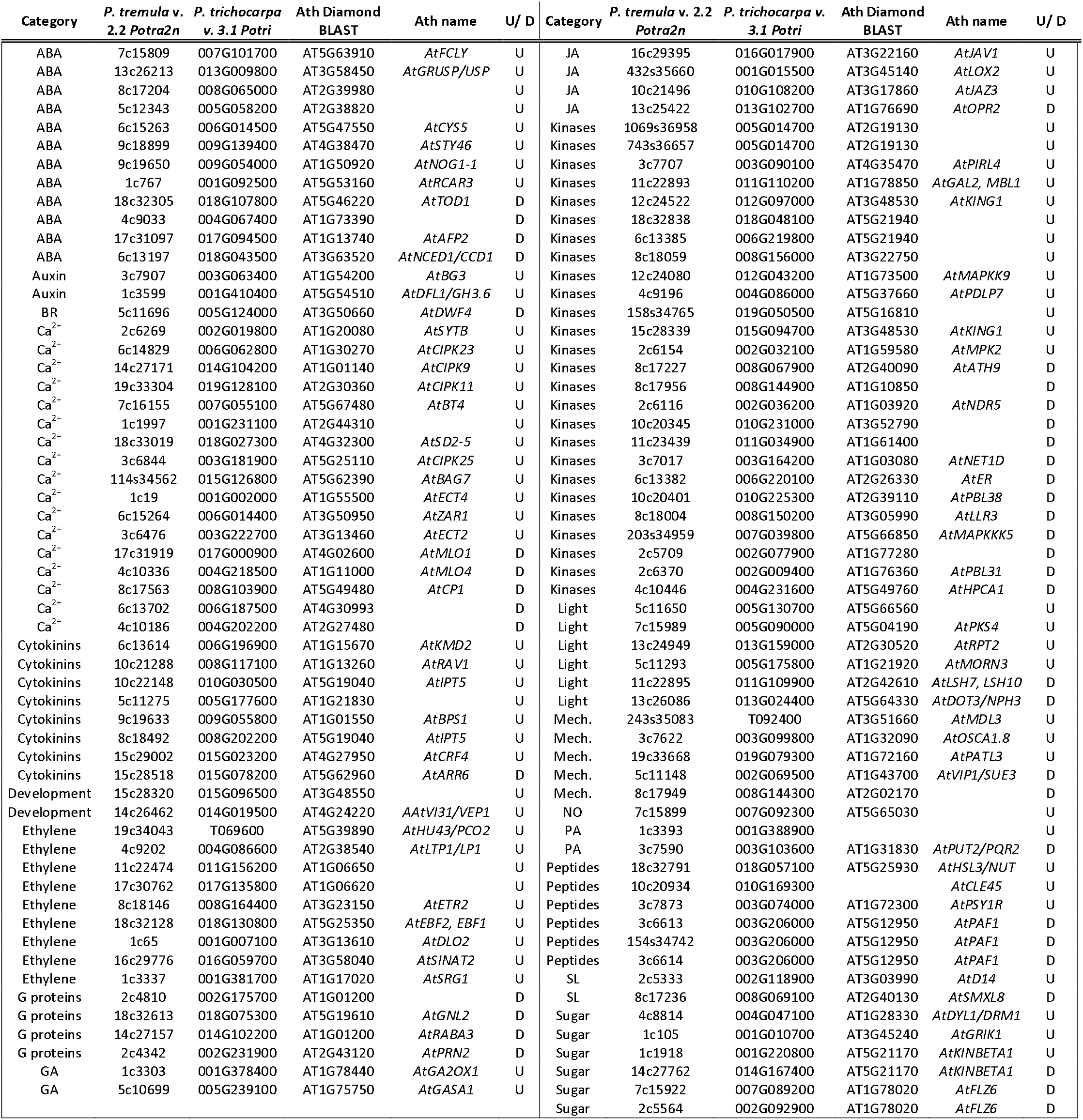
Signaling-related genes significantly up-(U) or downregulated (D) in common by GH10 and GH11 xylanases.

#### Transcriptomic changes in secondary wall-related genes

To find out if SW formation was affected by the xylanases at the transcript levels, we identified differentially regulated genes in transgenic lines with 35S:GH10, WP:GH10 and WP:GH11 constructs among known SW-related genes expressed in the wood forming tissues (**Supplementary Table S8**). Among cellulose-related genes, several genes from family GH9 encoding cellulases were found downregulated (**Table 2**). Among xylan-related genes, those involved in meGlcA substitution (*PtGUX1-A, PtGUX4-A* and *PtGXM1*), and acetylation (*PtXOAT1*, *PtRWA-A* and *PtRWA-B*) were found downregulated. Lignin biosynthesis pathway was also affected due to downregulation of genes involved in monolignol biosynthesis (*PtPAL4, PtCAld5H2,* homolog of *AtCAD6, Pt4CL3* and *5, PtCCoAOMT1* and *2*) and polymerization (*PtLAC12/AtLAC17*). This indicates that specific programs modifying cellulose, responsible for xylan substitution and lignin biosynthesis were downregulated. Among the master switches regulating these programs in *Populus* (Zhong et al., 2010; Ohtani et al., 2011), we identified two *VND6* genes *PtVND6-A2* and *PtVND6-C2* genes (named after Li et al., 2012 as listed in Takata et al., 2019) and their downstream TFs, *PtMYB199* homologous to *AtMYB85* - an activator of lignin biosynthesis, *PtSND2* and *PtNAC124* (homologous to *AtSND2*) and *PtMYB90*, *PtMYB161* homologous to *AtMYB52* activating cellulose and hemicellulose biosynthetic pathways (Zhong et al., 2008; Schuetz et al., 2013) downregulated in xylanase-expressing lines (**Table 2**). This suggests that specific sub-programs of SW biosynthesis have been downregulated via SW transcriptional cascade in transgenic lines. On the other hand, we also observed a strong downregulation of four AtMYB4 homologs, including PtLTF1that regulates lignification in response to stress (Gui et al., 2019), upregulation of PtMYB55 homologous to *AtMYB61* reported to positively regulate SW development in Arabidopsis coordinating a small network of downstream genes (Romano et al., 2012), and three homologs of AtMYB73 involved in salinity stress response and lateral root development (Kim et al., 2013; Wang et al., 2021).

**Table 2.**
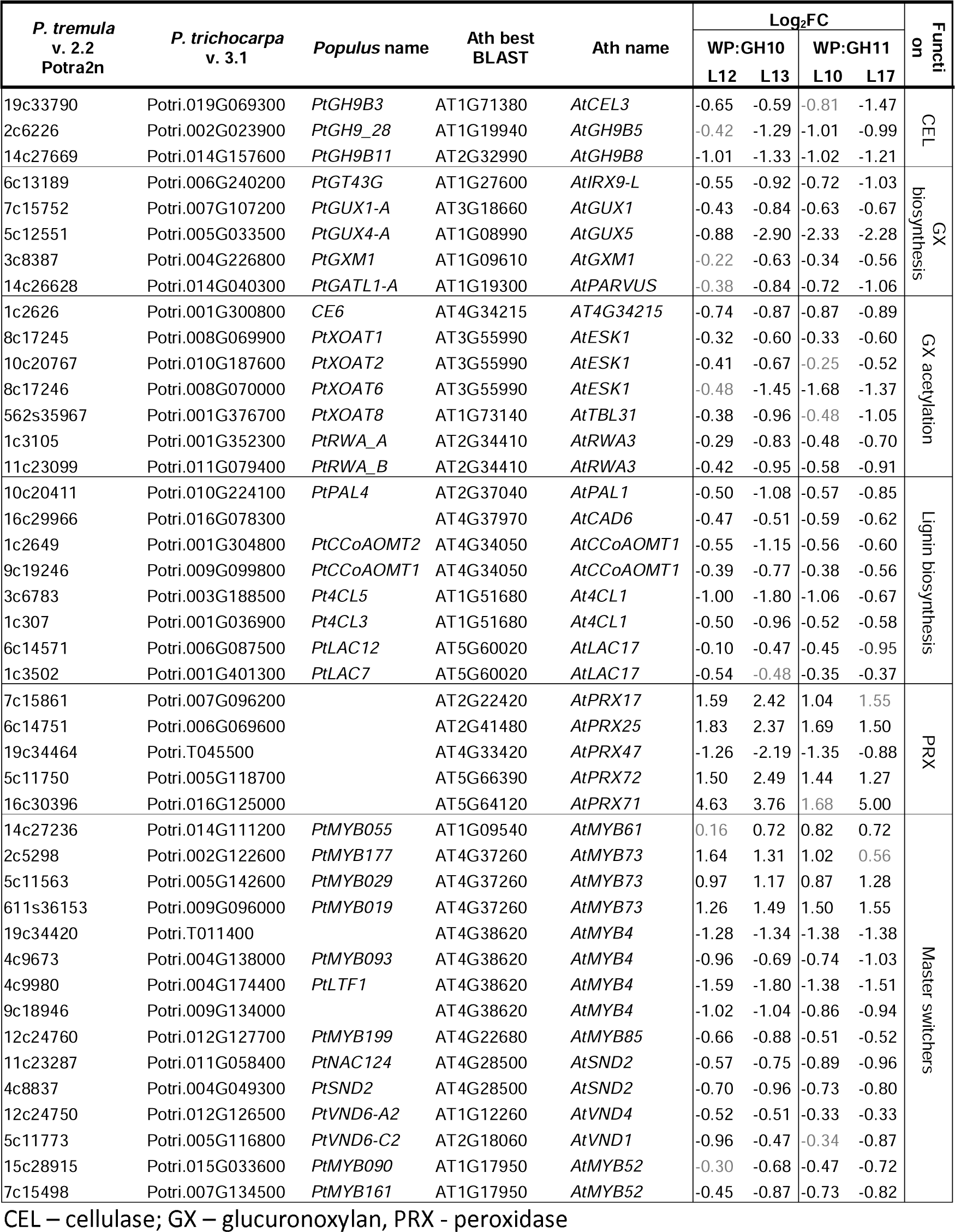
Cell wall-related genes significantly (P_adj_≤0.05) up- or downregulated in transgenic lines expressing GH10 and GH11 xylanases. Shown are values of Log_2_ fold change from WT levels. Values in grey are not significantly different from WT.

#### Co-expression networks formed by genes commonly affected in WP:GH10 and WP:GH11 lines

To identify co-expression networks of DEGs that might operate in developing wood we used AspWood database (Sundell et al., 2017). Eight networks were identified, the main network and seven side networks, and expression of the genes of each network in wood forming tissues, in different tree organs and in different xylanase-expressing lines was illustrated as heatmaps (**Fig. 10**, **Supplementary Figs S7-S10**, **Supplementary Table S7**). The main network included mostly genes expressed late during xylogenesis, but it contained smaller subnetworks of genes expressed during SW formation and during primary wall stage of xylem differentiation (**Supplementary Fig. S7**). It was dominated by signaling- and stress-related genes (**Fig. 10, Supplementary Table S7**), including many kinases, calcium signaling components, genes related to hormones (peptide - *PAF1s*, ethylene, ABA - *AtAFP2,* and *AtNCED1*, cytokinin - *AtARR6*), sugar responses, and stress (dehydration, salt, freezing and anoxia). The highly interconnected transcriptional factors included upregulated *AtWRKY75, AtERF110/PtERF57* and *AtLBD21/PtLBD047* and downregulated *AtLBD19*/*PtLBD043* and several *AtNAC074* homologs. The cell wall-related genes included downregulated *PtGH9B11, PtGH9_18, PtGUX4A*, and upregulated *AtXTH28/PtXTH40*.

**Figure 10.**
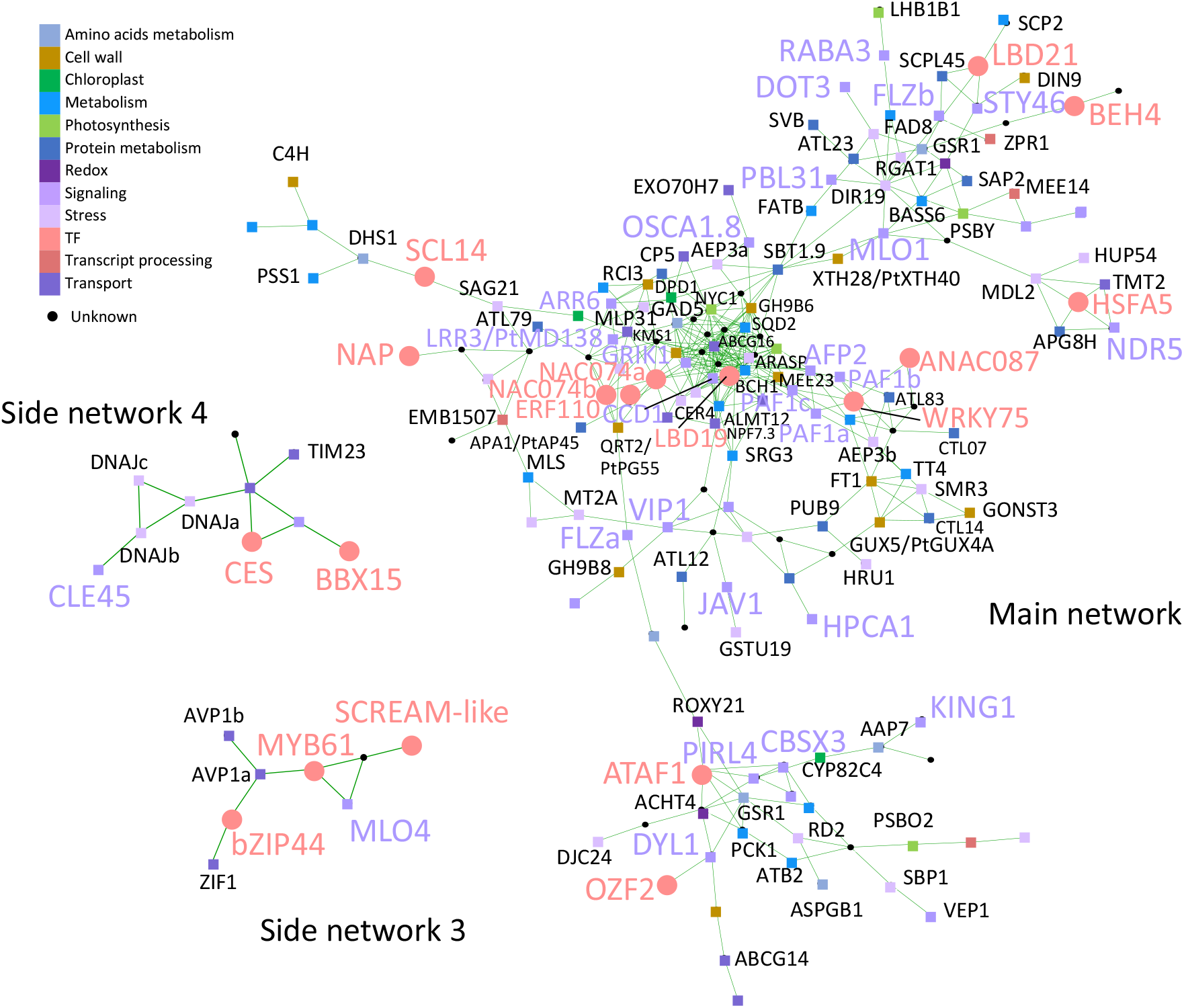
Co-expression networks of the core genes differentially expressed in GH10- and GH11-expressing aspen lines. Gene exclusive functional classification is indicated by the colors of the nodes, as listed in Table S7. Signaling- and stress-related genes and transcription factors are shown on large colored fonts. All gene names use *Arabidopsis* gene symbols, unless indicated by prefix Pt. For the gene expression data and all identified networks, please see Figures S7-S10.

Among the side networks, network 3 included genes differentially expressed in the cambium (**Fig. 10, Supplementary Fig. S9**), with three transcription factors, *AtMYB61, AtSCREAM-like* and *AtbZIP44,* the calcium-signaling related gene *AtMLO4* and transporters *AtZIF1* and *AtAVP1*. Side networks 4 and 6 were grouping genes expressed in the phloem that were almost all upregulated in transgenic lines (**Fig. 10, Supplementary Fig. S10**). Network 4 included *AtCLE45* encoding a peptide hormone and transcription factors *AtCES* and *AtBBX15*, whereas network 6 included a kinase (*AtMPK2*), peroxidase *AtPRX72* and stress-related genes (*AtGSTF11* and *AtKTI5*).

The side networks 1 and 2 (**Supplementary Fig. S9**) grouped genes highly upregulated by xylanases, expressed in the leaves, phloem and mature xylem zone, and involved in photosynthesis (*AtLHCA3, AtLHCB5, AtPSAD, AtPSAE, AtPSBR, AtPSBO2*) and photorespiration (*AtGOX1*).

Side networks 5 and 7 (**Supplementary Fig. S10 and 11**) grouped genes expressed at primary-SW transition in developing xylem and low expressed in extraxylary tissues, which were downregulated except one, *AtZPR1* encoding LITTLE ZIPPER1. Network 7 included *AtSTP10* encoding hexose-H^+^ proton symporter and *AtGNL2* encoding GNOM-like 2 an ARF guanidine exchange factor regulating vesicle trafficking.

#### Unique transcriptomic changes induced by the two families of xylanases

Venn diagrams (**Fig. 9A**) showed that 120 genes were jointly upregulated in two lines of WP:GH10 construct but were not affected in any of the lines of WP:GH11 construct whereas 230 genes were upregulated only in the GH11 expressing lines. Similar analysis of downregulated genes revealed 62 specifically downregulated in WP:GH10 and 117 in GH11 lines. Thus, GH11 altered expression of approx. two times more genes than GH10 and these genes were more frequently upregulated (**Supplementary Table S9**). GO analysis of these xylanase family-specific genes (**Supplementary Table 10)** revealed that GH11 induced more intense activation of stress and in particular raffinose stress-related genes than GH10.

## Discussion

### Fungal xylanases reduced xylan content and altered its structure in aspen secondary walls

The majority of transgenic lines expressing fungal GH10 and GH11 xylanases in cell walls had reduced TMS xylose content, and reduced molecular weight of SWE xylan. This was accompanied by reduced signals from LM10 antibodies in SWs and increased content of soluble xylose and xylobiose. Jointly these data demonstrate that the xylanases were active on cell wall xylan in aspen, cleaving the backbone. However, not all domains of xylan were equally susceptible to cell wall-targeted GH10 and GH11 xylanases. Analysis of degree of acetylation and OLIMP profiles after GH30 glucuronoxylanase digestion of SWE xylan indicated that highly acetylated and tightly glucuronidated regions of xylan were less prone to hydrolysis by these xylanases as these substitutions are known to restrict xylan backbone digestion by GH10 and GH11 xylanases (Biely et al., 2016; Kojima et al., 2022).

### Fungal xylanases affected wood cell wall structure and composition

*Irregular xylem phenotype* (*irx*) with thin-walled fibers and collapsed vessels is typical for mutants impaired in biosynthesis of SWs or one of its main components like cellulose, xylan and lignin (Jones et al., 2001; Turner et al., 2001; Brown et al., 2005; Persson et al., 2007; Hao et al., 2014). Here we show that *irx* phenotype was also induced by SW-targeted fungal xylanases. This SW thinning was without any change in cellulose content but with prominent decrease in lignin content and composition. While all lines had reduced content of G-lignin, only GH11-expressing lines had lower S-lignin content. Similarly, *Arabidopsis thaliana* expressing GH10 and GH11 xylanases had decreased lignin, especially G-lignin content in secondary xylem (Barbut et al., unpublished). Moreover, mutations in xylan biosynthetic genes were also observed to lead to the same lignin defects (Hao et al., 2014; Barbut et al., unpublished) suggesting an interdependence of lignification on xylan in SWs.

Whereas one mechanism of this interdependence has been proposed *via* xylan acting as a nucleation point for the polymerization of monolignols (Sapouna et al., 2023) our data point to a different possibility. First, we observed a severe decrease in mono- and oligolignols as well as phenolic glycosides suggesting severely decreased lignin monomer biosynthesis in xylanase-expressing lines. If the polymerization was affected *via* reduced lignin nucleation sites, one would expect increased content of unpolymerized lignols instead. Second, we observed downregulation of transcripts of lignification-specific *MYB*s and several genes involved in monolignol biosynthesis and polymerization. Similarly, key lignin biosynthetic genes were downregulated in *irx8/gaut12* mutant (Hao et al., 2014). Moreover, two negative regulators of phenylpropanoid biosynthesis encoding F-box proteins AtKFB and AtKMD2 targeting phenylalanine ammonia-lyase for ubiquitination (Zhang et al., 2013) were upregulated in xylanase-expressing lines (**Supplementary Table S7**). It is notewhorty that AtKMD2 was upregulated in xylobiose-treated Arabidopsis (Dewangan et al., 2023) and in ixr9 mutant (Faria-Blanc et al., 2018) that was also hypolignified (Petersen et al., 2012), making it a strong candidate for downregulation of phenylpropanoid pathway in xylan-compromised plants.

Xylanase-expressing aspen lines showed also downregulation of some key genes required for biosynthesis of MeGlcA and acetyl substitutions of GX. This could be a reaction counteracting the changes in GX substitution induced by xylanases in developing SW. This specific downregulation of GX biosynthesis subprogram likely involves specific branches of the SW regulatory program (Ohtani et al., 2011; Zhong et al., 2011; Taylor-Teeples et al., 2015; Chen et al., 2019), possibly including *VND6* homologs, the first layer master switches, and several members of the third layer master switchers. Some of these masters switchers, like *PtMYB161* (Wang et al., 2020a) of *PtLTF1* (Gui et al., 2019), could also participate in a feedback regulation of the SW biosynthesis in response to stress. An additional layer of suppression of SW program in transgenic plants could be mediated by decreased ABA levels since ABA signaling is needed to activate NST1 by phosphorylation (Liu et al., 2021). These observations are in line with our previous hypothesis based on observations in aspen with suppressed *PtGT43BC* expression that SW impairment is sensed by plants resulting in general shutdown of SW biosynthetic program (Ratke et al., 2018).

### Expression of fungal xylanases altered growth and vascular differentiation pattern in aspen

Modification of xylan backbone by SW-targeted GH10 and GH11 xylanases led to substantial decrease in height and biomass of trees. The previous experiments suppressing xylan backbone biosynthesis by knocking down *PtGT43A* and *B* (Lee et al., 2011), *PtGT43B* and *C* (Ratke et al., 2018), *PtGT47C (*Lee et al., 2009) or *PtGAUT12* (Li et al., 2011; Biswal et al., 2015) reported in contrast either no or positive effects on growth in *Populus,* especially when WP promoter was used. Intriguingly, the xylanases affected cambial growth by specifically inhibiting xylem formation and increasing phloem formation, which was correlated with increased cytokinin content. Phloem differentiation and cambial cell division are known to be regulated by local maxima in cytokinins which in turn exclude auxin maxima by regulating distribution of PIN transporters inhibiting auxin-dependent activation of HD-ZIPIII transcription factors and xylem differentiation (Bishopp et al., 2011; Immanen et al., 2016; Haas et al., 2022). Moreover, we found three homologs of *ZPR1* that is known to inactivate HD-ZIPIII transcription factors (Wenkel et al., 2007) upregulated in xylanase-expressing lines, which would provide additional mechanism suppressing xylem formation. Therefore, modification of xylan in SW appears to positively affect mitotic divisions in the cambium, enhance phloem differentiation, and in case of xylanase-expressing lines, inhibit xylem fate *via* transcriptional and hormonal regulation.

### Hormonal signaling pathways are affected in xylanase-expressing lines

Impairment of xylan integrity in SW by fungal xylanases induced severe systemic changes such as reduced plant height, reprogramming of cambial activity from xylem to phloem production, and suppression of SW formation program in differentiating xylem. Such changes require long- and short-distance signaling, which likely starts in differentiating xylem cells and involves plant hormones.

Decrease in ABA content and signaling was supported by a downregulation of a key ABA biosynthetic gene, homolog of *AtNCED1,* and the upregulation of homologs of negative regulators of ABA signaling pathway: *AtATAF1* (Garapati et al., 2015) and *AtKING1* (Papdi et al., 2008) in xylanase-expressing aspen. ABA forms a regulatory feedback loop with FERONIA (FER), a key RLK sensing cell wall integrity (Bacete and Hamann, 2020). ABA biosynthesis has been found to be downregulated after cell wall integrity signaling mediated by *At*THESEUS1 (*At*THE1) (Bacete et al., 2022), after stem mechanical disturbance (Urbancsok et al., 2023) and following *Botrytis cinerea* infection (Windram et al., 2012). On the other hand, ABA signaling was needed for increased biotic resistance in *Arabidopsis irx* mutants with defects in SW *CesA* genes (Hernández-Blanco et al., 2007).

Strigolactones (SLs) and/or related carotenoids have been previously shown to mediate *irx* phenotype and freezing tolerance of *esk1/tbl29* and other SW mutants impaired in cellulose and xylan biosynthesis in *Arabidopsis* (Ramírez and Pauly, 2019). In xylanase-expressing aspen lines the upregulation of a functional homolog of *AtDWARF14* (*AtD14*), *PtD14a,* encoding an SL receptor (Zheng et al., 2016) and *AtSMXL8* - involved in feedback regulation of SL signaling (Wang et al., 2020b)-supports activation of signaling by SLs. Moreover, a downregulation of a chalcone synthase transcript *AtTT4* which controls flavonoid biosynthesis downstream SLs (Richmond et al., 2022) was observed in common in xylanase-expressing aspen and xylobiose-treated *Arabidopsis* (Dewangan et al., 2023). Xylanases also induced *BYPASS1* (*BPS1*) encoding a plant-specific inhibitor of a carotene-related xylem-transported hormone inhibiting shoot development (Van Norman and Sieburth, 2007).

Xylanases also affected ethylene signaling as evidenced by increased ACC levels, and upregulation of several ethylene related genes including *ETHYLENE RESPONSE FACTORS* (*ERF*s) which were also induced in xylobiose-treated *Arabidopsis* (Dewangan et al., 2023), and a homolog of *ERF1* regulating growth under stress (Hoang et al., 2020). Upregulated JA signaling was also evident based on transcriptome analysis. This signaling pathway has been implicated in cell wall integrity response downstream of THE1 (Bacete et al., 2022). Both ethylene and JA signaling pathways were stimulated in the developing xylem by mechanical stress (Urbancsok et al., 2023).

Upregulation of cytokinins in xylanase-expressing aspen expectantly would increase plastid multiplication resulting in strong upregulation of photosynthesis-related genes, and lipid and amino acid metabolism. Transcriptomics data supported these hypotheses with upregulation of a homolog of *AtPLASTID DIVISION2* (*AtPDV2)* that regulates plastid division (Chang et al., 2017) and many genes involved in plastid organization and photosynthesis. Among several cytokinin-related DEGs, a homolog of *AtARR6* encoding a negative regulator of cytokinin response was downregulated. *ARR6* has been implicated in cell wall modification and immunity (Bacete et al., 2020).

Thus, xylan integrity impairment caused by xylanases affected signaling *via* ABA, strigolactones/carotenes, ethylene and cytokinins, which overlaps with primary cell wall integrity signaling, and responses to mechanical and other abiotic and biotic stresses (Bacete and Hamann, 2020; Rivero et al., 2021).

### Local candidates for stress perception in secondary wall-forming cells

The perception of xylan impairment in SW expectedly would involve local sensors including xylobiose (DAMP) sensors (Dewangan et al., 2023) and other cell wall integrity sensing components (Bacete and Hamann, 2020). One of them could be *HPCA1* encoding a novel RLK responsible for H_2_O_2_ perception at the plasma membrane and activation of calcium influx (Wu et al., 2020). Several other calcium signaling-related genes were upregulated, including mechanosensitive calcium channel *AtOSCA1.8* (Yuan et al., 2014; Murthy et al., 2018), defense-activated calcium channel *AtZAR1* (Bi et al., 2021), *AtMDL3* known to be dependent on activity of mechanically activated MCA channels (Mori et al., 2018), *AtLBD38* regulated by calcium influx via cyclic nucleotide-gated channel CNG15 (Tipper et al., 2023). Furthermore, *AtMLO4* and *AtMLO1* homologs were downregulated. *AtMLO4* is a calcium channel involved in mechanical stress and gravitropism signaling (Zhu et al., 2021). Among candidates expressed at primary-SW transition other transporters were also identified, including *PtVP1.1* (*AtAVP1*) encoding a pyrophosphate-fueled proton pump regulating apoplastic pH and involved in stress responses (Yang et al., 2015), *AtSTP10* encoding a proton-coupled sugar symporter responsible for uptake of monosaccharides from apoplast into plant cells (Bavnhøj et al., 2021), and *AtGNL2* involved in ER-Golgi trafficking of proteins (Teh and Moore, 2007). Some of these genes were regulated in common with xylobiose-treated *Arabidopsis* (Dewangan et al., 2023) (**Supplementary Table S7**).

### Xylanase-induced changes in cell wall chemistry improved wood saccharification potential

Xylan binds to cellulose surfaces and interconnects lignin and was shown to impede the enzymatic saccharification (De Martini et al., 2013). Moreover, as it is the main source of yeast-inhibiting acetic acid, it is predicted to inhibit the fermentation (Donev et al., 2018). Therefore, decreasing xylan content and its modification are considered as effective strategies for improving biomass biorefinery properties. Here, we show that expressing either GH10 or GH11 xylanases in aspen SWs greatly improved glucose yield and production rate per wood weight in saccharification without pretreatment which were doubled or even tripled compared to WT. Previous experiments with *HvXyl1* expressed in poplar reported a 50% increase in glucose yield in saccharification after steam pretreatment (Kaida et al., 2009). Even milder xylan reduction by suppressing *GT43* genes of clades B and C resulted in increased in glucose yield in saccharification without pretreatment by 30% to 40%, but a negligible effect was observed after acid pre-treatment (Lee et al., 2011; Ratke et al., 2018). The high glucose yields observed in the present study were however associated with growth penalties. On the other hand, no such penalties were observed in *GT43*-suppressed aspen either in the greenhouse or in the field (Ratke et al., 2015; Derba-Maceluch et al., 2023). It is therefore evident that saccharification benefits and growth are not necessarily negatively linked. Our current analysis of transcriptomic and metabolomic changes revealed many candidates for uncoupling regulation of growth and development from xylan reduction in xylanase-expressing lines. Elucidation of their function could lead to designing better strategies to obtain saccharification-improved plants that grow just as well as WT or even better.

### Conclusions

This study evaluated the effects of postsynthetic modification of xylan backbone by overexpression of fungal xylanases on growth, secondary cell wall characteristics and wood properties in aspen. Our results demonstrated that xylanases decreased the content of xylan and its molecular weight, and modified its substitution pattern. This inhibited tree growth, wood production, SW development and lignin biosynthesis. The hormonomics, metabolomics and transcriptomics analyses revealed that xylan impairment activated hormonal signaling and affected genetic regulatory pathways that modified cambial growth and adjusted SW biosynthesis program, suggesting the activation of SW integrity sensing. Although the benefits of highly enhanced glucose yield in saccharification from transgenic wood biomass were offset by growth penalty, the identified candidates for the SW integrity sensing mechanism could be used to uncouple beneficial and undesirable effects for developing improved lignocellulose in aspen for biorefinery.

## Experimental Procedures

### Generation of transgenic lines

The *Aspergillus nidulans* cDNA clones encoding GH10 (AN1818.2; GenBank: ABF50851.1) and GH11 (ANIA_03613; NCBI_GeneID:2873037, XP_661217.1) xylanases (Bauer et al., 2006) were used to generate expression vectors. The signal peptide of GH10 was replaced by the hybrid aspen (*Populus tremula* L. x *tremuloides* Michx.) signal peptide from gene *PtxtCel9B3* (alias *PttCel9B*) (GenBank AY660968.1; Rudsander et al., 2003) as described previously (Gandla et al., 2015), whereas native fungal signal peptide was used for GH11 vector. The cloning primers are listed in **Supplementary Table S11**. The entry clones generated using the pENTR/D-TOPO cloning system (Thermo Fisher Scientific, Uppsala, Sweden) were used to make the expression clones in either pK2WG7.0 (Karimi et al., 2002) for ectopic expression using 35S promoter or in pK-pGT43B-GW7 (Ratke et al., 2015) for expression specifically in cells developing secondary cell walls driven by the wood-specific promoter (WP). The resulting vectors (35S:GH10, 35S:GH11, WP:GH10 and WP:GH11) were introduced into competent *Agrobacterium tumefaciens* (Smith and Townsend, 1907) Conn 1942, strain GV3101 using electroporation. Binary vectors were transformed into hybrid aspen (*Populus tremula* L. × *tremuloides* Michx., clone T89) as described previously (Derba-Maceluch et al., 2015). Lines with the highest transgene expression were selected from 20 independent lines for further analyses.

### Plant growth in the greenhouse

*In vitro* propagated saplings were planted in soil (K-jord, Hasselfors Garden AB, Örebro, Sweden) in 7 L plastic pots, watered to 25% - 30% (v:v) soil moisture content, covered with transparent 8 L plastic bags, and grown for nine weeks in the phenotyping platform (WIWAM Conveyor, custom designed by SMO, Eeklo, Belgium) as described by Wang et al. (2022) under 18 h /6 h (day/night) light regime with 160-230 µmol m^-2^ s ^-1^ light intensity during the day, 22 °C /18 °C temperature, and the average air relative humidity of 60%. White light (FL300 LED Sunlight v1.1) and far-red light (FL100 LED custom-made, 725-735 nm) lamps from Senmatic A/S (Søndersø, Denmark) were used for illumination. After two weeks the bags were removed, and plants were watered automatically based on weight, their height was automatically measured.

At the end of experiment, trees were photographed, and stems diameters at base and aboveground fresh weights were recorded. A 30 cm-long stem segment above internode 37 was debarked, frozen in liquid nitrogen and stored at −70 °C for RNA, metabolomics and hormonomics analyses. The stem below was used for determining internode length. The 38^th^ and 39^th^ internodes were used for microscopy analyses. The four-cm long bottom segment was used for SilviScan analysis, and the remaining stem was debarked and freeze-dried for 48 h for wood chemistry analyses. Belowground biomass was determined by weighing cleaned and air-dried roots.

### Wood microscopy analysis

For light microscopy, samples of three trees per line were fixed in FAA (4% formaldehyde, 5% acetic acid, 50% ethanol). Transverse sections (40-50 µm-thick) were prepared with a vibratome (Leica VT1000S, Leica Biosystems, Nussloch, Germany) and stained with safranin-alcian blue (Urbancsok et al., 2023). Lignin autofluorescence was analyzed at 470 nm (Kitin et al., 2020). Images were acquired by Leica DMi8 inverted microscope (Leica Biosystems, Germany) equipped with digital camera and analyzed with ImageJ software.

Another set of samples from the same trees were fixed in 0.1% glutaraldehyde, 4% paraformaldehyde, 50 mM sodium cacodylate buffer for 4 h at room temperature and embedded in LR white resin as described elsewhere (Pramod et al., 2014). Two µm-thick sections were cut using an ultramicrotome (RMC Powertome XL, USA) and stained with toluidine blue O for light microscopy analysis. Transverse ultrathin sections (70-90 nm-thick) were prepared using an ultramicrotome Ultracut E (Leica Biosystems) with a diamond knife and mounted on copper grids. For lignin localization, sections were stained with KMnO_4_ (Donaldson, 1992). The xylan immunogold labelling with LM10 monoclonal antibody was carried out as described by Pramod et al. (2014). All sections were examined with a transmission electron microscope (FEI TALOS L120C) at an accelerating voltage of 100 kV. Cell wall thickness and gold particle density was determined using ImageJ based on ten and twenty measurements per tree, respectively, for two lines per construct.

### SilviScan analyses

A SilviScan instrument (RISE, Stockholm, Sweden) was used for determining wood and fiber properties of six trees per line, 24 per WT as described by Urbancsok et al. (2023).

### Cell wall chemical analyses

For initial pyrolysis and TMS analyses, wood powder from three trees per line was obtained by filing the freeze-dried wood and sieving the sawdust with Retsch AS 200 analytical sieve shaker (Retsch GmbH, Haan, Germany) to 50-100 µm.

Py-GC/MS assay used 50 µg (± 10 µg) of powder in a pyrolyser equipped with autosampler (PY-2020iD and AS-1020E, Frontier Lab, Japan) connected to a GC/MS (7890A/5975C, Agilent Technologies Inc., Santa Clara, CA, USA). The pyrolysate was processed and analyzed according to Gerber et al. (2012).

Alcohol-insoluble residue (AIR) was prepared as described by Gandla et al. (2015). AIR was destarched by α-amylase (from pig pancreas, cat. nr. 10102814001, Roche, USA) and amyloglucosidase (from *A. niger* cat. nr.10102857001, Roche) enzymes and the matrix sugar composition was analyzed by methanolysis-trimethylsilyl (TMS) procedure as described by Pramod et al. (2021). The silylated monosaccharides were separated by GC/MS (7890A/5975C; Agilent Technologies Inc., Santa Clara, CA, USA) according to Gandla et al. (2015). Raw data MS files from GC/MS analysis were converted to CDF format in Agilent Chemstation Data Analysis (v.E.02.00.493) and exported to R software (v.3.0.2). 4-*O*-Methylglucuronic acid was identified according to Chong et al. (2013). The amount of monosaccharide units per destarched AIR weight was calculated assuming their polymeric form.

For the remaining cell wall analyses, the pith was removed from debarked and freeze-dried stem segments the segments of seven trees per line were ground together using Retsch Ultra Centrifugal Mill ZM 200 (Retsch GmbH, Haan, Germany) equipped with a 0.5 mm ring sieve. The resulting wood powder was then sieved by Retsch AS 200 vibratory sieve shaker to isolate powder with particle size of 50-100 and 100-500 µm.

The 50-100 µm fraction was used in triplicates for monosaccharide analysis by a two-step sulfuric acid hydrolysis (Saeman et al., 1954). In brief, 1 mg of sample was incubated with 125 μL of 72% H_2_SO_4_ at room temperature for 3 h, then diluted with 1375 μL of deionized water and incubated at 100°C for 3 h. Hydrolyzates were diluted 10 times with MilliQ water, filtered through 0.2 mm syringe filter (Chromacol 17-SF-02-N) into HPAEC-PAD vials and analyzed by high performance anion exchange chromatography with pulsed amperometric detection (HPAEC-PAD) (ICS-6000 DC, Dionex) equipped with a CarboPac PA1 column (4 × 250 mm, Dionex) at 30°C using the eluent gradients previously reported (McKee et al., 2016). Quantification of monosaccharides was performed by standard calibration of ten monosaccharides (Ara, Rha, Fuc, Xyl, Man, Gal, Glc, GalA, MeGlcA and GlcA) with concentrations between 0.005 and 0.1 g L^-1^.

For subcritical water extraction, 2 g of 100-500 µm wood powder was extracted with 0.2 M formate buffer, pH 5.0, at 170 °C and 100 bar in an accelerated solvent extractor (ASE-300, Dionex, USA). Extraction proceeded in 4 steps with residence times of 10, 20, 30 and 60 min according to Sivan et al., (2023). Low-molecular weight compounds were removed by dialysis using Spectra/Por 3 membranes (Spectrum, USA), and the extracted polymers were freeze-dried.

For alkaline extraction, 1 g of wood powder with particle size 100-500 µm was incubated with 24% KOH for 24 h at room temperature (Escalante et al., 2012; Timell, 1961), filtered through 60 µm wire mesh and neutralized with 0.4 vol of acetic acid. Hemicellulose was precipitated with 96% ethanol (4 °C for overnight), centrifuged, washed in 80% ethanol, dissolved in distilled water and freeze-dried. Molar mass of extracts was determined by size exclusion chromatography coupled to refractive index and UV-detectors (SECurity 1260, Polymer Standard Services, Mainz, Germany). The samples (2 mg) were dissolved in 1 mL of dimethyl sulfoxide (DMSO Anhydrous, Sigma-Aldrich) with 0.5% w/w LiBr (Anhydrous free-flowing Redi-Dri, Sigma-Aldrich) at 60 °C, and filtered through 0.45 µm PTFE syringe filters (VWR). The separation was carried through GRAM Analytical columns of 100 and 10000 Å (Polymer Standard Services, Mainz, Germany) at a flow rate of 0.5 mL min^-1^ and 60 °C. The columns were calibrated using pullulan standards between 345 and 708 000 Da (Polymer Standard Services, Mainz, Germany).

The acetyl content of water extracts was determined in duplicates by overnight saponification of approx. 5 mg of sample in 1.2 mL of 0.8 M NaOH at 60 °C with constant mixing, neutralization with 90 µL of 37% HCl and filtration through 0.45 mm Chromacol syringe filters (17-SF-02(N), Thermo Fisher Scientific). The released acetic acid was detected by UV at 210 nm using high pressure liquid chromatography with UV detector (Dionex-Thermofisher Ultimate 3100, USA) and separation by a Rezex ROA-organic acid column (300 x 7.8 mm, Phenomenex, USA) at 50 °C in 2.5 mM H_2_SO_4_ at 0.5 mL/min. Propionic acid was used as an internal standard.

For oligosaccharide mass profiling (OLIMP), the alkaline and 30 min water extracts were digested using GH10 endo-β-(1-4)-xylanase from *Cellvibrio mixtus* (Megazyme), a GH11 endo-1,4-β-xylanase from *Neocallimastix patriciarum* (Megazyme) and GH30 endo-1,4-β glucuronoxylanase (kindly provided by Prof. James F. Preston, University of Florida), incubating 1 mg of extract in 1 mL of 20 mM sodium acetate buffer (pH 5.5) and 10 U enzyme for 16 h at 37 °C. After enzyme inactivation at 95°C for 10 min, the hydrolysates were ten times diluted in acetonitrile 50 % (v/v) with 0.1 % (v/v) formic acid and filtered through Chromacol 0.2 μm filters (Scantec Nordic, Sweden). Samples were then briefly passed through a ZORBAX Eclipse Plus C18 column 1.8 μm (2.1 × 50 mm) (Agilent Technologies, Santa Clara, CA) and the oligosaccharide profiles were analyzed by HPAEC-PAD as reported previously (McKee et al., 2016) using xylooligosaccharides (X_2_-X_6_; Megazyme) as external standards and electrospray ionization mass spectrometry (ESI-MS) with a Synapt HDMS mass spectrometer (Waters, USA) in positive-ion mode and capillary and cone voltage set to 3 kV and 70 kV, respectively. The oligosaccharides were detected as [M + Na]+ adducts.

Oligosaccharide sequencing was achieved after the separation of labeled oligosaccharides by tandem LC-ESI-MS/MS. Derivatization was performed by reductive amination with anthranilic acid as previously described (Mischnick, 2012). The labelled oligosaccharides were separated through an ACQUITY UPLC HSS T3 column (150 × 2.1 mm, Waters, USA) at a flow rate of 0.3 mL min^-1^ and a gradient of increasing acetonitrile content (10–30%) over 40 min. Mass spectrometric analysis was performed in positive mode with the capillary voltage and cone set to 3 kV and 70 kV, respectively. MS2 was performed by selecting the ion of interest [M + Na]+ through single ion monitoring and subjecting it to collision-induced dissociation using argon as the collision gas, at a ramped voltage of 35–85 V. Assignment of proposed structures was performed by reference to labeled standards and analysis of the fragmentation spectra using ChemDraw (PerkinElmer, Waltham, Massachusetts, USA).

### Saccharification assay

Three technical replicates from each line and six from WT were used for analytical-scale saccharification. Wood powder moisture content was measured using Mettler Toledo HG63 moisture analyzer (Columbus, OH, USA) and 50 mg of dry material was used per sample. Acid pretreatment was carried out using an Initiator single-mode microwave instrument (Biotage Sweden AB, Uppsala, Sweden) with 1% (w/w) sulfuric acid at 165 °C for 10 min. Enzymatic hydrolysis without or after acid pretreatment was performed at 45 °C using 4 mg of the liquid enzyme mixture Cellic CTec2 (cat. nr. SAE0020, Sigma-Aldrich, Saint Louis, MO, USA) as previously described (Gandla et al., 2021). Samples were analyzed for glucose production rate at 2 h by using an Accu-Chek^®^ Aviva glucometer (Roche Diagnostics Scandinavia AB, Solna, Sweden) following the calibration with a set of glucose standard solutions. After 72 h, the yields of monosaccharides were quantified using HPAEC-PAD (Ion Chromatography System ICS-5000 by Dionex, Sunnyvale, CA, USA) (Wang et al., 2018).

### RNA analyses

Developing xylem tissues were scrapped from the debarked frozen stem and ground in a mortar with a pestle in liquid nitrogen. Approximately 100 mg of fine tissue powder was extracted with CTAB/chloroform:isoamylalcohol (24:1) followed by LiCl and sodium acetate/ethanol precipitation to isolate total RNA (Chang et al., 1993).

RNA samples from three trees per line were DNase treated with DNA-free^TM^ kit (cat. nr. AM1906, Thermo Fisher Scientific, Waltham, MA, USA) then reverse-transcribed using iScript^TM^ cDNA synthesis kit (cat. nr. 1708891) (Bio-Rad Laboratories, Hercules, CA, USA) following the manufacturers’ instructions. Quantitative polymerase chain reactions (qPCRs) were performed using LIGHTCYCLER 480 SYBR GREEN I Master Mix (Roche, Indianapolis, IN, USA) in a Bio-Rad CFX384 Touch Real-Time PCR Detection System with 10 µL reaction volume. PCR program was 95°C for 3 min, then 50 cycles of 95°C for 10 s, 55°C for 10 s and 72°C for 15 s. UBQ-L (Potri.005G198700) and ACT11 (Potri.006G192700) were selected as reference genes from four tested genes based on GeNorm (Vandesompele et al., 2002). The primer sequences are listed in **Supplementary Table S11**. The relative expression level was calculated according to Pfaffl (2001)

For transcriptomics, RNA was purified as described previously (Urbancsok et al., 2023) and four or five biological replicates per transgenic line and eight biological replicates of the WT with RNA integrity number (RIN) ≥ 8 were used for cDNA preparation and sequencing using NovaSeq 6000 PE150 at Novogene Co., Ltd. (Cambridge, United Kingdom). Quality control and mapping to the *P. tremula* transcriptome (v.2.2), retrieved from the PlantGenIE resource (https://plantgenie.org; Sundell et al., 2015) were carried out by Novogene. Raw counts were used for differential expression analysis in R (v3.4.0) with the Bioconductor (v.3.4) DESeq2 package (v.1.16.1), as previously detailed (Kumar et al., 2019).The best BLAST hits were identified in *Populus trichocarpa* (v3.1) and *Arabidopsis thaliana* (v11.0).

### Hormonomics and metabolomics

Frozen developing xylem samples were ground as described above. Hormone profiling was done according to Šimura et al. (2018), with slight modifications (Urbancsok et al., 2023). ACC (1-aminocyclopropane-1-carboxylic acid) was quantified according to Karady et al. (2024).

Metabolites were extracted and analyzed as described by Abreu et al. (2020) and Urbancsok et al. (2023) and processed by an untargeted approach. The generated data were normalized against the internal standard and weight of each sample. Changes in abundance between transgenic and WT samples were considered as significant if P[≤[0.05 (t-test) and |fold change|[≥[1.5. The false discovery rate was <[0.05.

### Statistical analyses

Unless otherwise stated, statistical analyses were performed in JMP Pro (v.16.0) software (SAS Institute Inc., Cary, NC, USA).

## Acknowledgements

This work was supported by the Knut and Alice Wallenberg (KAW) Foundation, the Swedish Governmental Agency for Innovation Systems (VINNOVA), Swedish Research Council, Kempestiftelserna, T4F, Bio4Energy and the SSF program ValueTree RBP14-0011 to EJM. MK was supported by The Czech Science Foundation (GAČR) via 20-25948Y junior grant.

We are grateful to the undergraduate student Rakhesh Vaasan, KTH, for help with cell wall analyses. We acknowledge the Umeå Plant Science Centre (UPSC) Tree Phenotyping Platform, Bioinformatics Facility, Microscopy Facility, Biopolymer Analytical Platform, and the Swedish Metabolomics Centre in Umeå.

## Author Contributions

PS performed majority of wood chemistry and microscopy analyses, interpreted the data and wrote the manuscript. JU processed wood material, extracted RNA, analyzed transgene expression and prepared samples for omics analyses. JU, END and FRB carried the greenhouse experiment and tree phenotyping. END performed bioinformatic analyses. MDM created transgenic aspen and collected samples for Silviscan analysis. JŠ, KC and MK analysed hormones. MLG and LJJ analysed saccharification potential. MM analyzed metabolomics data. EH and FV carried out hemicellulose analyses. ERM designed cloning strategy. EJM designed and coordinated the research, secured the funding, and finalized the paper with contributions from all authors.

## Data availability

The raw RNA-Seq data that support the findings of this study are available in the European Nucleotide Archive (ENA) at EMBL-EBI (https://www.ebi.ac.uk/ena/browser/home), under accession no. PRJEB61635 and…..

