## Supplementary Figures S1-S10 for "Modification of xylan in secondary walls alters cell wall biosynthesis and wood formation programs"

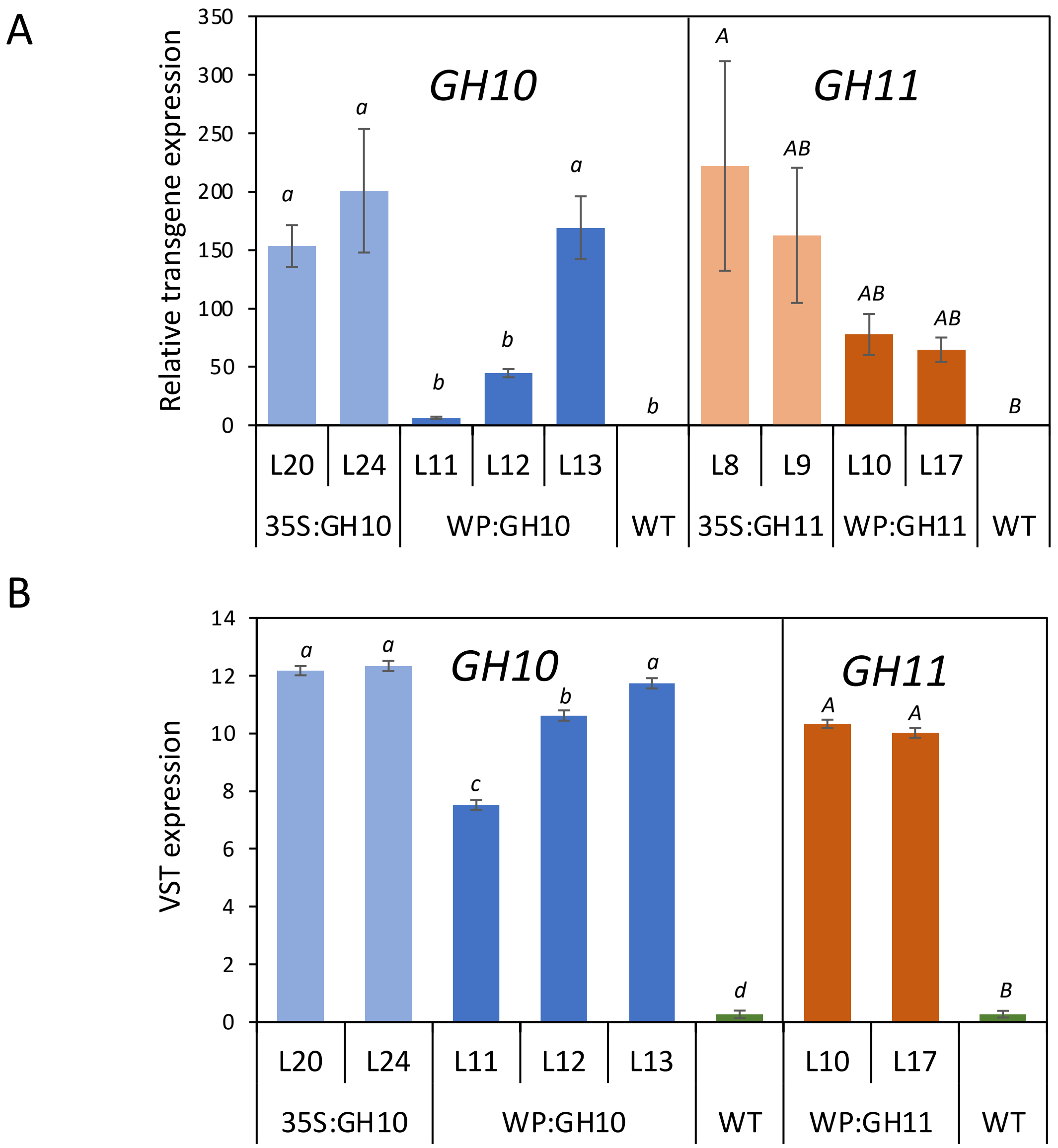


**Supplementary Figure S1. Transgene expression levels in developing wood of transgenic lines expressing GH10 and GH11 xylanases. (A)** Expression by RT-PCR using actin and ubiquitin genes for calibration and normalized to the lowest-expressing transgenic line. WT expression represents the noise. **(B)** Transcript quantitation based on RNA sequencing. Data are means ± SE, N = 3 for transgenic lines and 6 for WT in (A) or N=5 for transgenic lines and 8 for WT in (B). Different letters above the bars in B indicate significant difference among averages (P≤0.05, Tukey test).

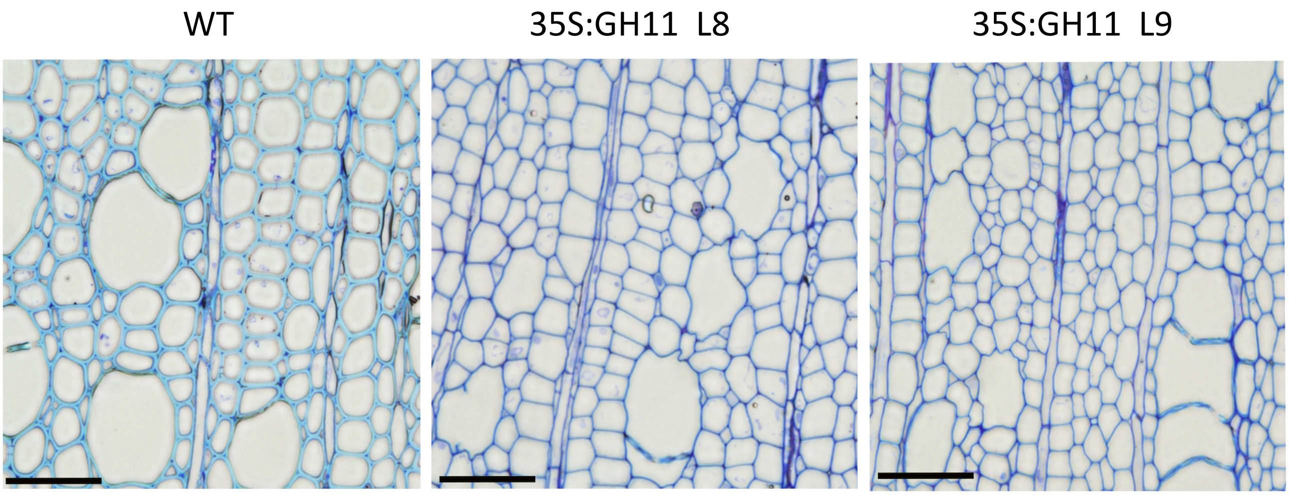


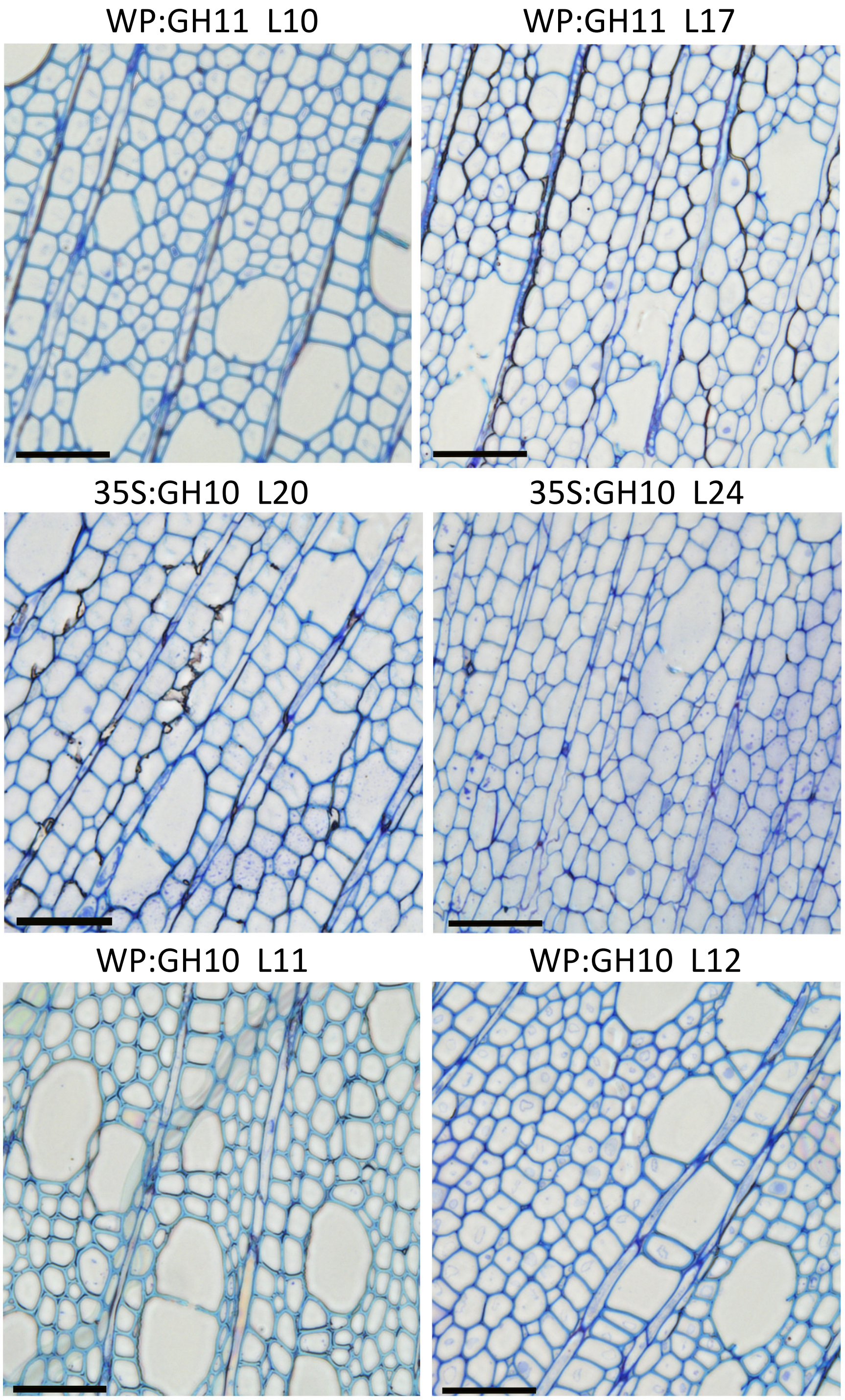


**Supplementary Figure S2. Toluidine blue stained wood sections showing reduction in cell wall thickness and change in staining indicative of reduced lignin content in transgenic lines expressing xylanases.** Some vessel elements in most affected lines show *irregular xylem phenotype* (*irx*). The phenotype is visible in all analyzed lines except WP:GH10_line 11, which had lower transgene expression than other lines. Scale bar = 50 µm

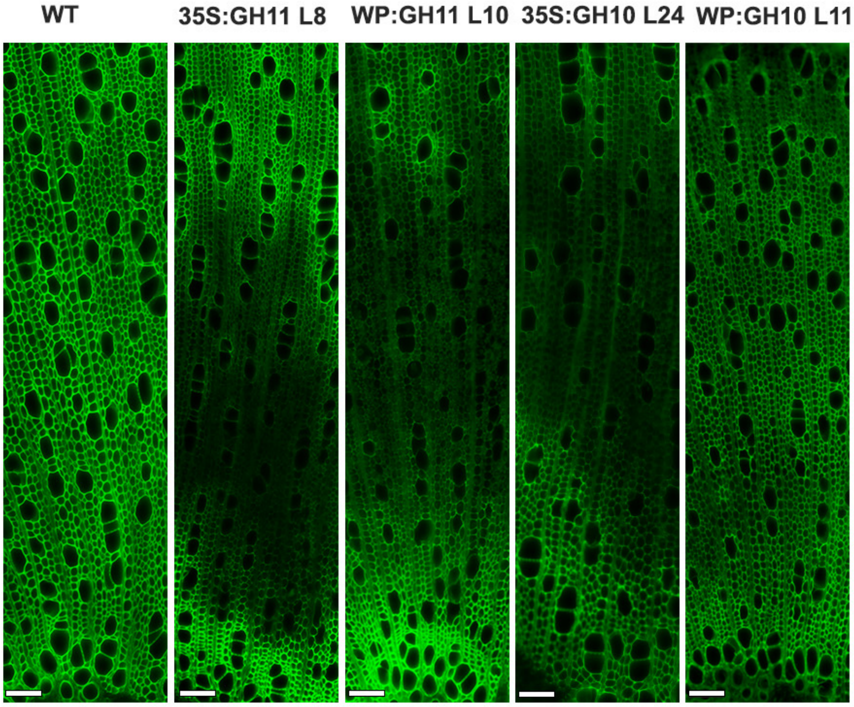


**Supplementary Figure S3. Fluorescence microscopy for detection of lignin in the wood tissue of transgenic lines expressing GH10 and GH11 xylanases.** Note the weak autofluorescence from the irregular xylem phenotypes of transgenic line. Scale bar= 50µm.

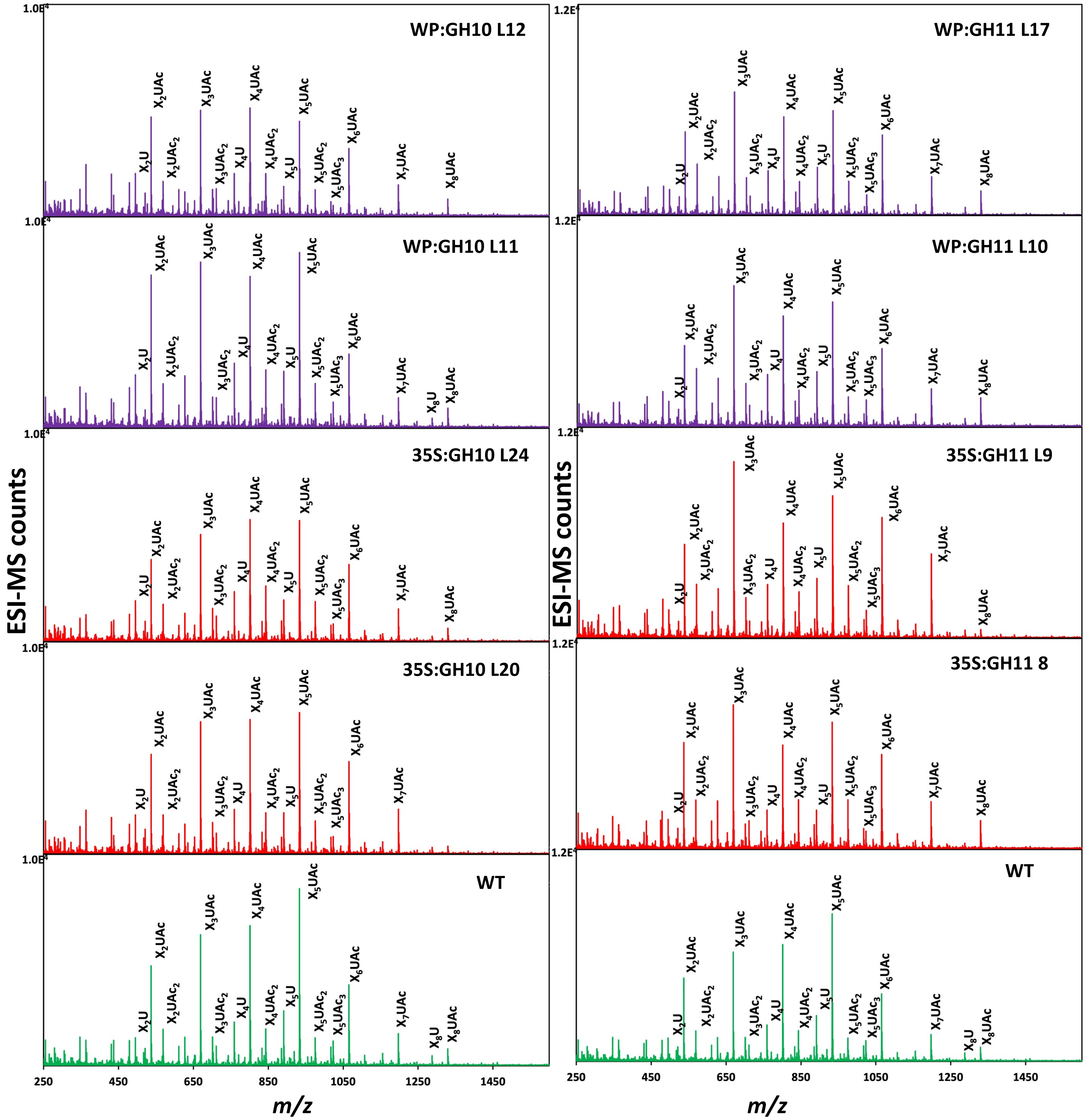


**Supplementary Figure S4. Oligomeric mass profiling (ESI-MS) of acetylated glucuronoxylan extracted with 30 min subcritical water extraction from transgenic lines expressing GH10 and GH11 xylanases. The oligomers are released by incubating the extracted hemicellulose with GH30 glucuronoxylanase.**

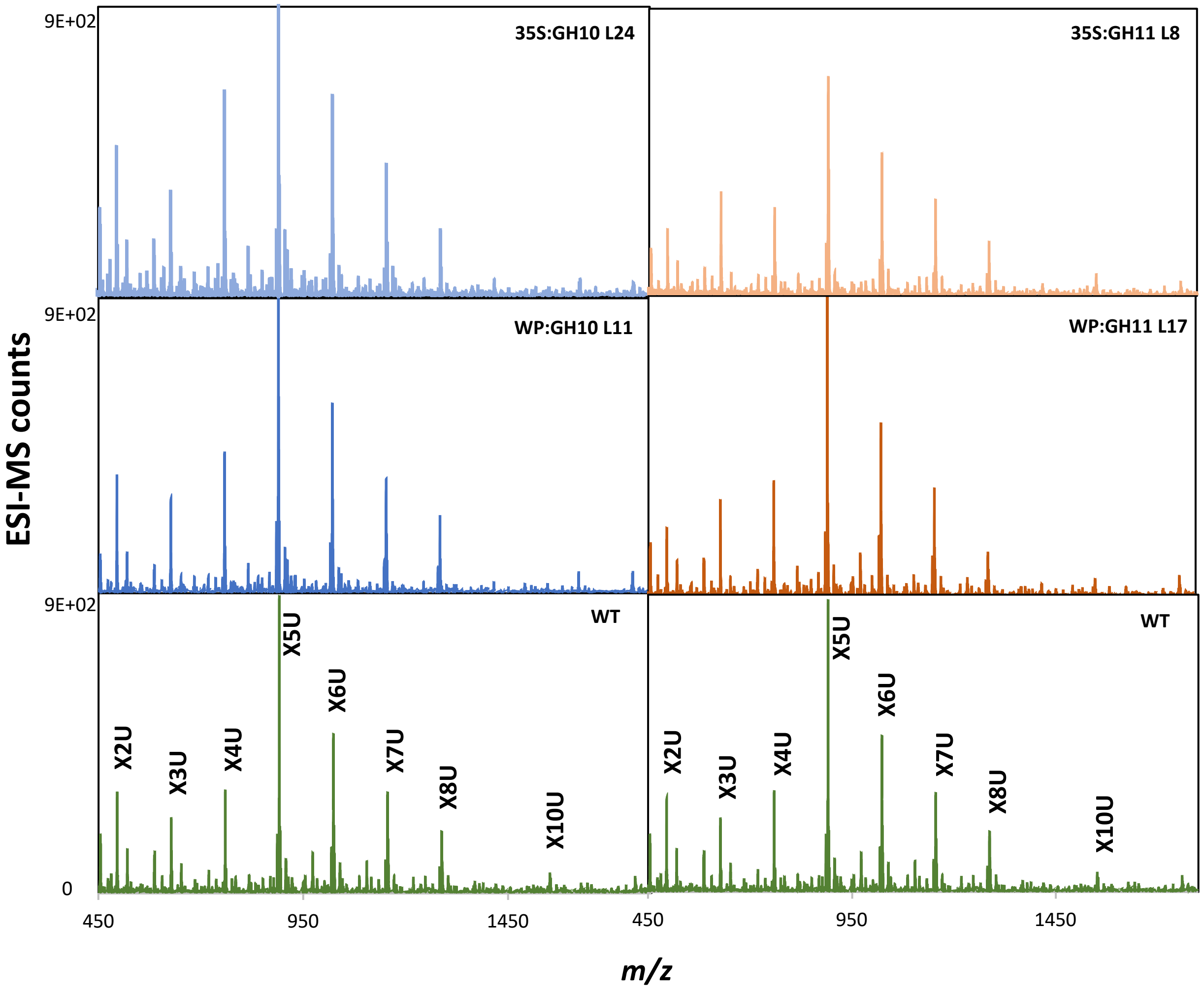


**Supplementary Figure S5. Oligomeric mass profiling (ESI-MS) of glucuronoxylan extracted with alkali from transgenic lines expressing GH10 and GH11 xylanases. The oligomers are released by incubating the extracted hemicellullose with GH30 glucuronoxylanase.**

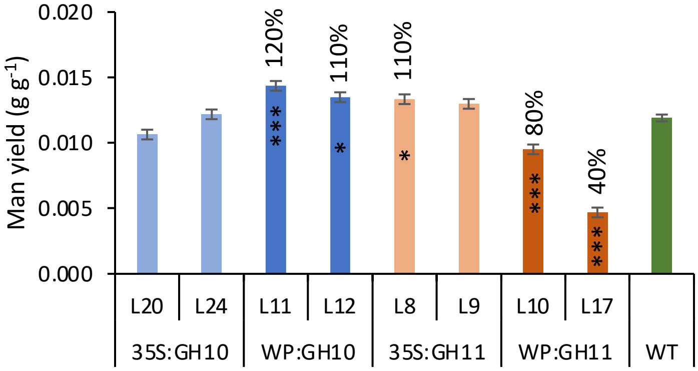


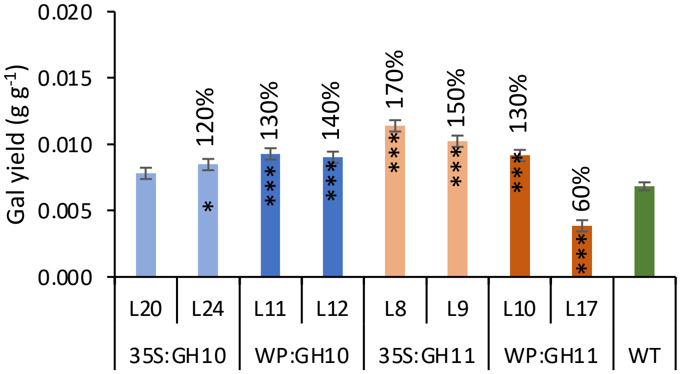


**Supplementary Figure S6. Saccharification yields of mannose and galactose obtained from wood of transgenic lines expressing GH10 and GH11 xylanases.** The sugars were released during acid pretreatment. Data are means ± SE, N = 3 or 6 technical replicates from the pooled material of 6 trees for transgenic lines and WT, respectively. * - P≤0.05; ** - P≤0.01; *** - P≤0.001 for comparisons with WT by Dunnett's test.

**Figure S7. Main co-expression network of the core genes differentially expressed in both lines expressing GH10 and both lines expressing GH11 xylanases in the wood-forming tissues and their expression patterns in different tissues and transgenic lines. (A)** Main co-expression network in the wood-forming tissues colored according to gene expression in wood developmental zones shown in the heatmap **(C). (B)-(D)** Heatmaps showing expression of the genes from the main expression network **(A)** in different tissues of aspen **(B),** wood developmental zones **(C),** and in transgenic lines (Log$_2$ fold change compared to wild type) **(D).**

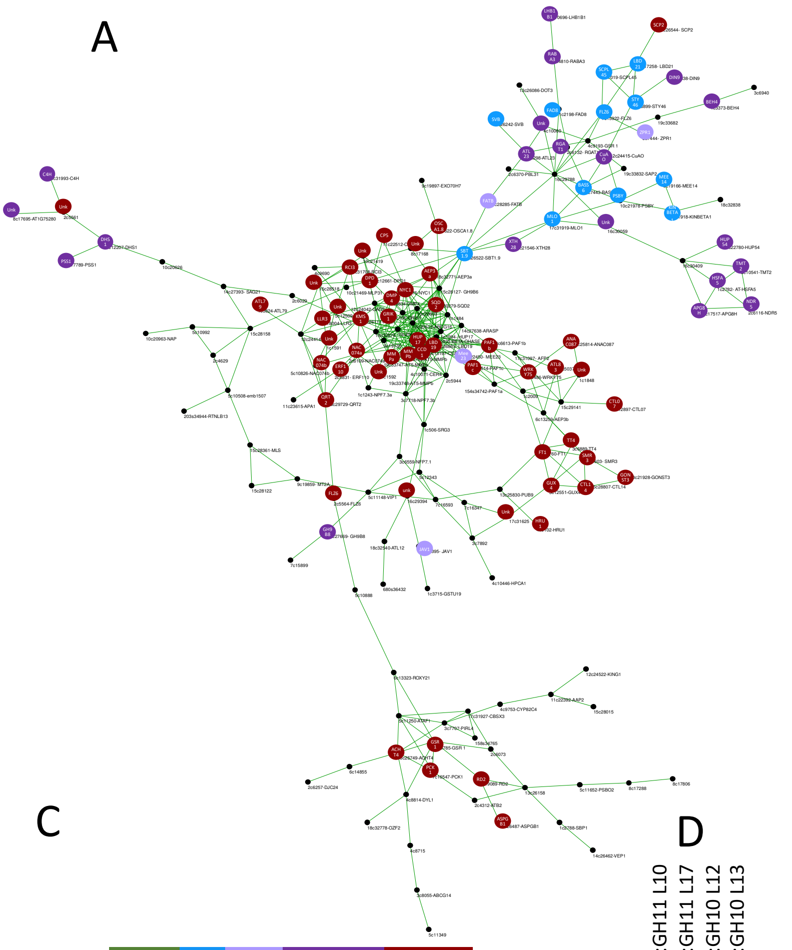


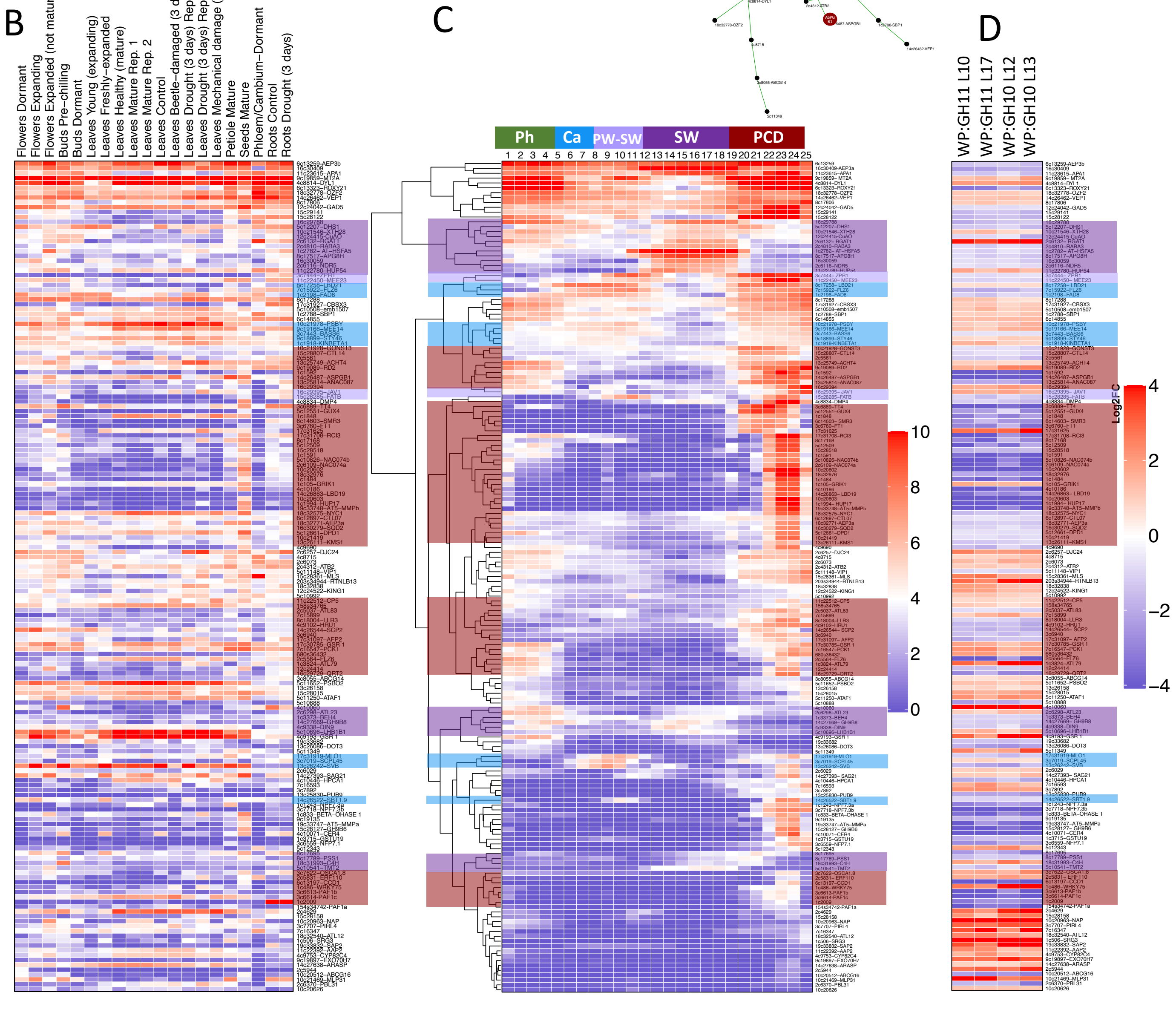

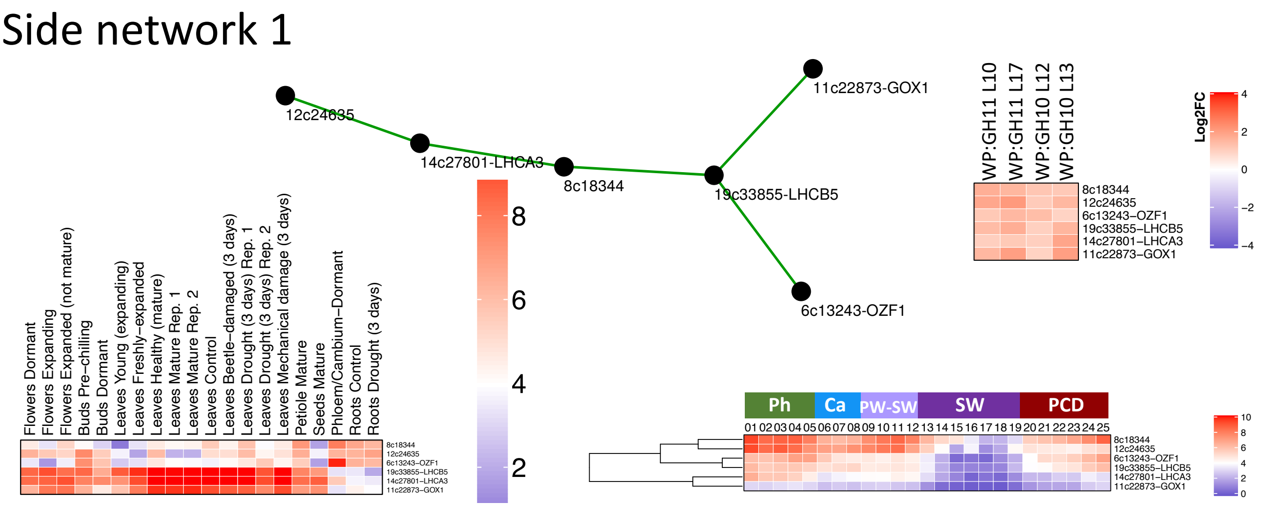


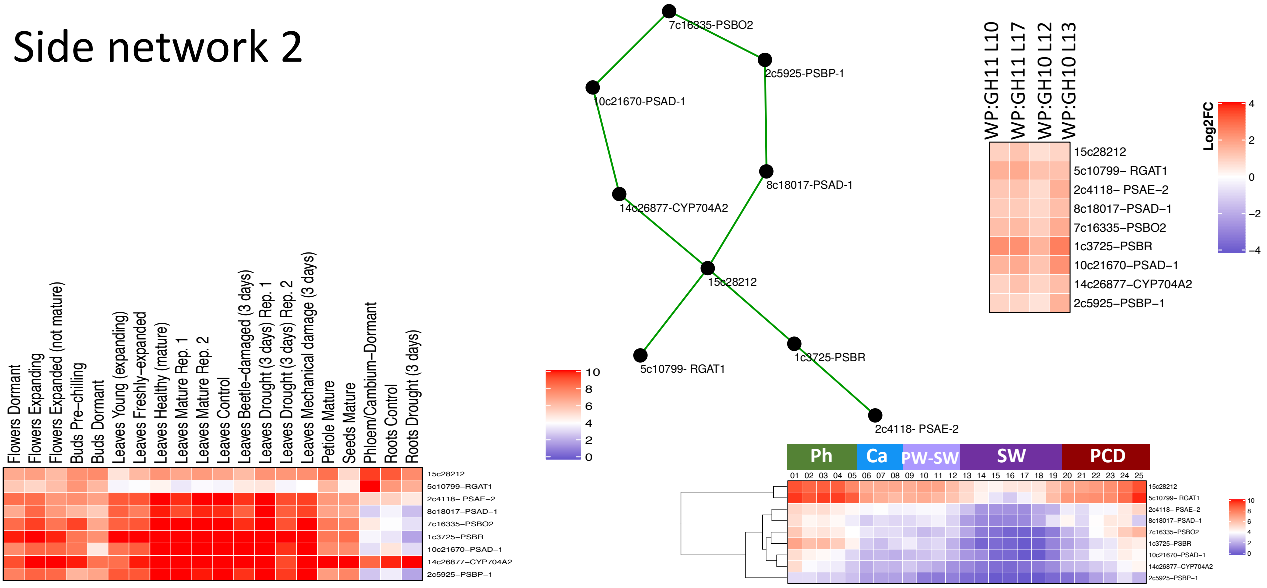


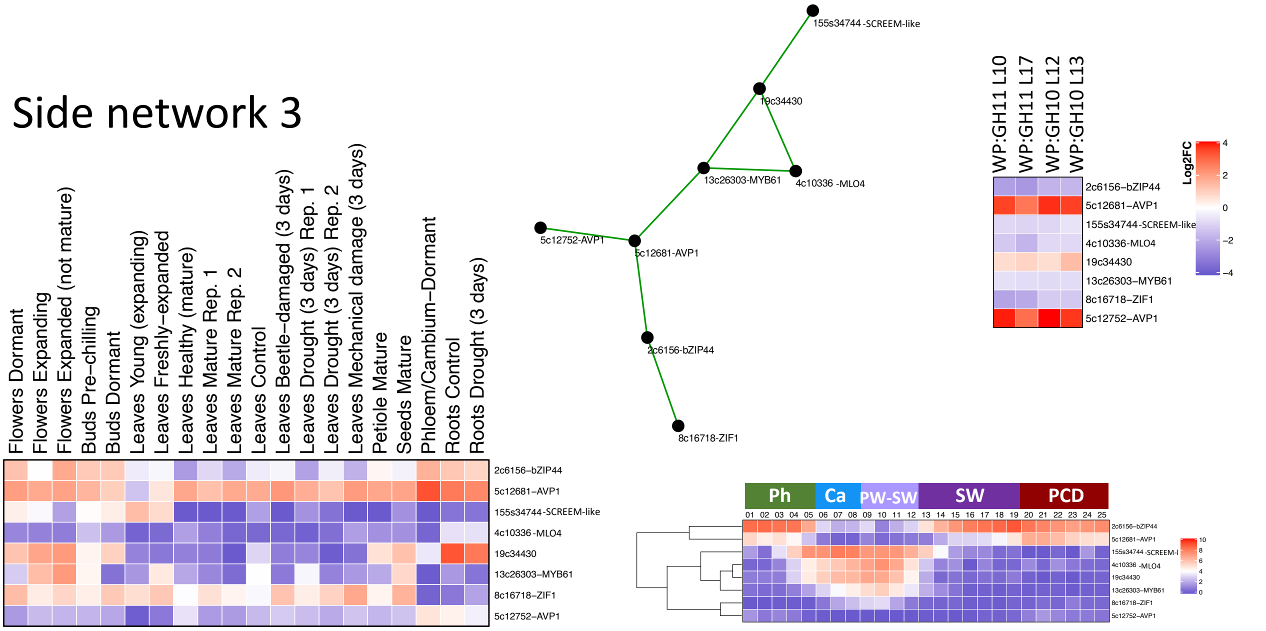


**Figure S8. Side co-expression networks 1-3 of the core genes differentially expressed in both lines expressing GH10 and both lines expressing GH11 xylanases in the wood-forming tissues and their expression patterns in different tissues and in transgenic lines.** Note that network 3 includes genes downregulated or upregulated in the cambium.

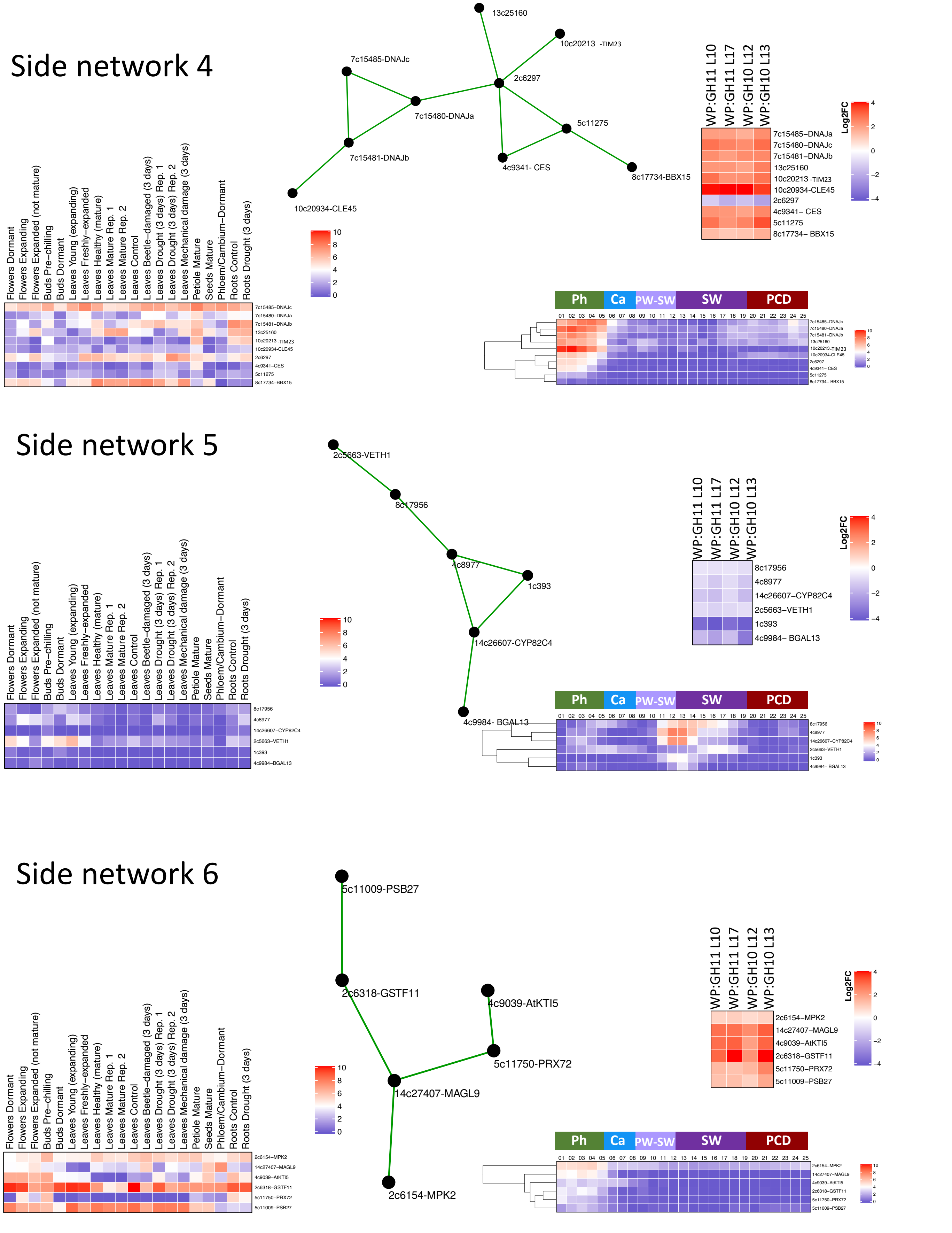


**Figure S9. Side co-expression networks 4-6 of the core genes differentially expressed in both lines expressing GH10 and both lines expressing GH11 xylanases in the wood-forming tissues and their expression patterns in different tissues and in transgenic lines.** Note that networks 4 and 6 include genes specifically expressed in the phloem, which are mostly upregulated in transgenic lines, whereas network 5 includes genes upregulated during primary to secondary wall transition, which are downregulated in transgenic lines.

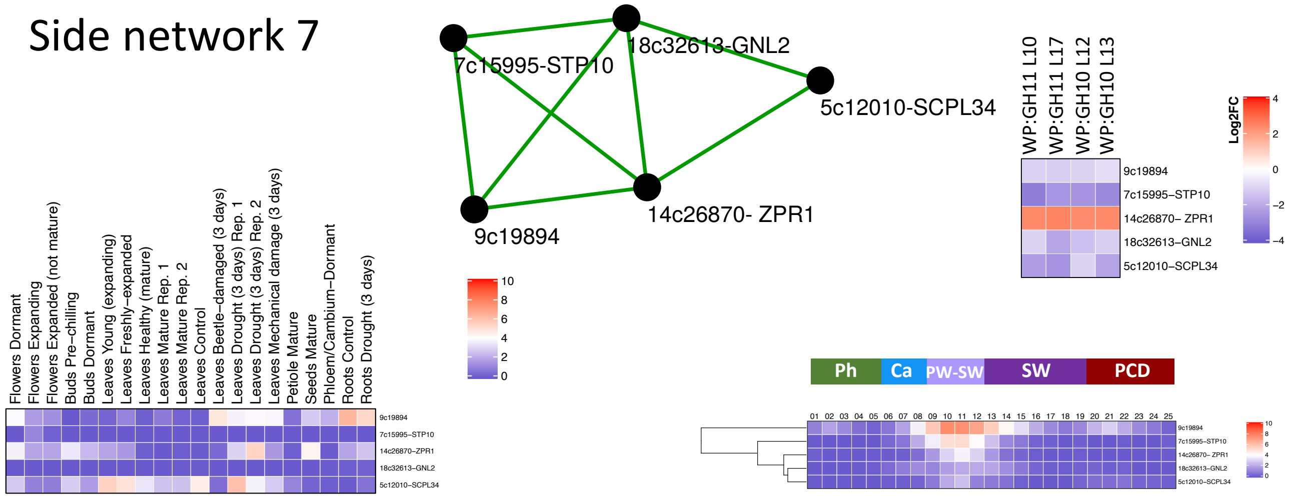


**Figure S10. Side co-expression network 7 of the core genes differentially expressed in both lines expressing GH10 and both lines expressing GH11 xylanases in the wood-forming tissues and their expression patterns in different tissues and in transgenic lines.** Note that network 7 includes genes specifically upregulated during primary to secondary wall transition, which are downregulated in transgenic lines except for one, ZPR1.
